# Distinct RopGEFs successively drive polarization and outgrowth of root hairs

**DOI:** 10.1101/534545

**Authors:** Philipp Denninger, Anna Reichelt, Vanessa A. F. Schmidt, Dietmar G. Mehlhorn, Lisa Y. Asseck, Claire E. Stanley, Nana F. Keinath, Jan-Felix Evers, Christopher Grefen, Guido Grossmann

## Abstract

Root hairs are tubular protrusions of the root epidermis that significantly enlarge the exploitable soil volume in the rhizosphere. Trichoblasts, the cell type responsible for root hair formation, switch from cell elongation to tip growth through polarization of the growth machinery to a pre-defined root hair initiation domain (RHID) at the plasma membrane. The emergence of this polar domain resembles the establishment of cell polarity in other eukaryotic systems [1–3]. Rho-type GTPases of plants (ROPs) are among the first molecular determinants of the RHID [4, 5] and later play a central role in polar growth [6]. Numerous studies have elucidated mechanisms that position the RHID in the cell [7–9] or regulate ROP activity [10–18]. The molecular players that target ROPs to the RHID and initiate outgrowth, however, have not been identified. We dissected the timing of the growth machinery assembly in polarizing hair cells and found that positioning of molecular players and outgrowth are temporally separate processes that are each controlled by specific ROP guanine nucleotide exchange factor (GEFs). A functional analysis of trichoblast-specific GEFs revealed GEF3 to be required for normal ROP polarization and thus efficient root hair emergence, while GEF4 predominantly regulates subsequent tip growth. Ectopic expression of GEF3 induced the formation of spatially confined, ROP-recruiting domains in other cell types, demonstrating the role of GEF3 to serve as a membrane landmark during cell polarization.

## RESULTS AND DISCUSSION

### Temporal analysis of hair cell polarization reveals phased deployment of the tip-growth machinery

To dissect the process of breaking cellular symmetry and initiating polar growth in plants, we analyzed the gradual assembly of the tip growth machinery in trichoblasts using specific markers for cytoskeletal rearrangement, plasma membrane specialization, cell wall modification, vesicle trafficking, and Rho-GTPase signaling. Using stable transgenic *Arabidopsis* lines expressing fluorescently labeled versions of these protein markers, we determined the timing of their polarization at the RHID during trichoblast differentiation. As the *Arabidopsis* root represents a time-axis of development from stem cells to mature cells, we numbered the developmental stages (−7 to +3) within a cell file, with the last cell before bulging labeled −1 and the first cell after the onset of bulging labeled +1 (**Figure 1A**). We used the integral plasma membrane marker GFP-LTI6B [19] as a reference and calculated the polarity index of fluorescently labeled marker proteins to quantify protein accumulation at the RHID (**Figure 1B, C**). Our survey unveiled a two-phase assembly of the tip growth machinery: an initiation phase, during which the RHID is positioned and predefined, followed by the tip growth phase. Consistent with previous reports [4, 5], mCitrine-labeled (mCit) GTPase ROP2 associated with the RHID during the early initiation phase (stage −4), well before any detectable cell bulging (**Figure 1B-D, Figure S1**). The timing of mCit-ROP2 polarization appeared to be largely independent from promoter activity, as analysis of an estradiol-inducible reporter construct for mCit-ROP2 resulted in a similar timing of RHID association, albeit with higher cytosolic signal (**Figure 1D, Figure S1**). Closely related GTPases ROP4 and ROP6 exhibited a similar timing as ROP2, when inducibly expressed (**Figure 1D, Figure S1**), which indicates that ROP GTPases are prepositioned well before growth is activated. The only tested ROP-effectors that were found to polarize after ROPs but still during the initiation phase, were the PHOSPHATIDYLINOSITOL-4-PHOSPHATE 5-KINASE 3 (PIP5K3) [20], and the NADPH oxidase ROOT HAIR DEFECTIVE 2 (RHD2) [21, 22]. PIP5K3 and RHD2 have both been shown to directly interact with ROPs during tip growth regulation [18, 23] and modulate vesicle traffic to the plasma membrane and cell wall rigidity, respectively [24].

**Figure 1.**
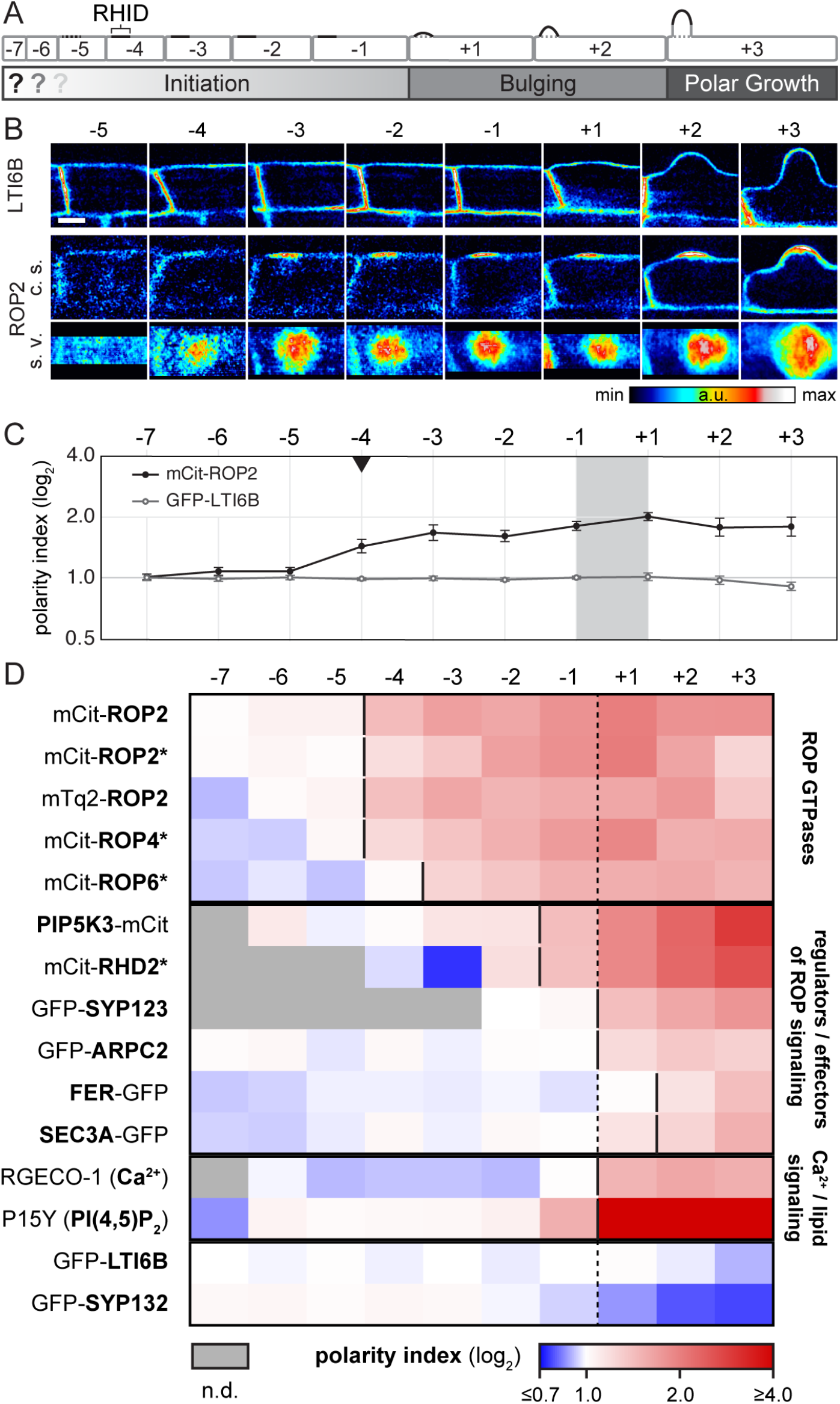
Timeline analysis of protein accumulation at the RHID. (**A**) Model of root hair development in an epidermal cell file. During the initiation phase the root hair initiation domain (RHID) emerges in elongating trichoblasts and the tip growth machinery is assembled using an unknown mechanism. Stages −1/+1 mark the stages before and after the onset of bulging followed by the transition into polar growth. (**B**) Comparison of mCit-ROP2 and GFP-LTI6B protein localization at the RHID in different stages of root hair development. For mCit-ROP2, cross sections (c. s.) and surface views (s. v.) of the same cells are shown. Intensities are represented in false-color, covering the full range of measured values within each data set. Scale bar, 10 μm. (**C**) Polarity index values (signal intensity in/out of RHID; log_2_ scale) for mCit-ROP2 and GFP-LTI6B along the different root hair stages. Error bars indicate SEM (n(stage −1)=11(ROP2); 14(LTI6B)). The black triangle indicates the first stage with a significant difference between mCit-ROP2 and the evenly distributed GFP-LTI6B (*p* < 0.01). (D) Protein accumulation at the RHID throughout the early development of root hairs in 13 different marker lines. Genes are sorted by the timing of accumulation within their group of protein function. Color coding represents the polarity index for each developmental step. (n.d.: no data). For each stage average polarity index values from multiple cells are shown (see **Figure S1** for details). Stages −7 to −1 represent cells in the initiation phase before bulging, +1 to +3 cells after bulging. Solid vertical lines indicate the first stage with a significant difference between the respective marker and GFP-LTI6B. Listed genes were fused to either eGFP or mCitrine (mCit), under control of their own promoters, except for UBQ10::P15Y (PI(4,5)P_2_ reporter), 35S::LTI6B and genes under control of an estradiol inducible promoter (indicated by an asterisk). See also **Figure S1**.

The second phase of root hair emergence was characterized by the polarization of known ROP effectors such as (ARPC2), a subunit of the actin polymerizing ARP2/3 complex [25], and the markers for vesicle transport and secretion SYP123 and SEC3A [26–28] (**Figure 1D, Figure S1**). The onset of bulge formation was further characterized by local elevation of cytosolic Ca^2+^ as reported by the sensor RGECO-1 [29, 30], as well as the formation of PI(4,5)P_2_ at the emerging apex as indicated by the reporter P15Y [31]. The receptor-like kinase FERONIA, which activates ROPs during root hair tip growth [32], accumulated after bulging in stage +2 (**Figure 1D, Figure S1**). Even though our visualization of total ROP distribution does not provide information on the ROP activity state, the temporal separation of ROP positioning and growth activation led us to hypothesize that distinct upstream, phase-specific regulators drive ROP GTPase recruitment and activation.

### RopGEFs exhibit specific spatiotemporal patterns of expression and subcellular localization

ROPs have been extensively studied during cell elongation [33], stomata regulation [34], pavement cell development [35, 36], and in regulation of the two tip-growing plant cell types, pollen tubes [37] and root hairs [4, 5]. As molecular “switches”, the activity-state of GTPases depends on regulators that “toggle” the switch between active or inactive [38, 39]. GTPase-activating proteins (GAPs) accelerate hydrolysis of a bound GTP to GDP, rendering the GTPase inactive. Examples for GAPs with described roles in pollen tube growth are RopGAP1 [40] and REN1 [14]. To switch the GTPase into an active state, guanyl-nucleotide exchange factors (GEFs) facilitate GDP release from the GTPase and the binding of another GTP molecule, and thereby promote interactions with downstream effectors [39, 41]. RopGEFs of the PRONE family [42, 43] and SPK1, the only DOCK family GEF [44], play known roles in tip growth regulation in pollen tubes [17, 45, 46] and leaf trichomes[44, 47]. However, temporal relationships between ROPs and RopGEFs have not yet been established in these cell types and much less is known about RopGEF functions in root hairs.

To identify candidate ROP regulators that play potential roles in root hair initiation and growth regulation we mined cell-type-specific gene expression data [48–50] and compared expression profiles of RopGEFs in root trichoblasts and atrichoblasts. Among these RopGEFs (hereafter abbreviated as GEFs), the expression of GEF3 and GEF4 showed the strongest root hair-specificity, followed by GEF12, GEF11, GEF14, and GEF10 in descending order (**Figure S2A**). Based on their trichoblast-specific expression, these GEFs were selected for subsequent analyses. To determine in which stage of hair cell maturation the candidate GEFs play their roles, we analyzed their expression patterns in growing roots expressing mCit fusions under the control of the respective native promoters. We observed a diverse distribution of expression patterns among the remaining GEFs, with a surprisingly unique pattern for each gene (**Figure 2A**). We further analyzed the timing of polarization at the RHID for each mCit-labeled GEF and compared it to the timing of mCit-ROP2. We found that mCit-GEF3 and mCit-GEF14 reached significant polarization consistently one cell stage earlier than mCit-ROP2 (**Figure 2B, Figure S1**). A major difference between both expression patterns was that GEF14 expression lasted from the meristematic zone (MZ) to the elongation zone (EZ), whereas GEF3 expression was first observed in the root transition zone (TZ, in-between MZ and EZ) and lasted beyond hair emergence (**Figure 2A**). The beginning of gene expression of GEF4 and GEF12 was detected first in EZ and the polarization of both proteins at the RHID coincided with the onset of bulge formation (**Figure 2A, B, Figure S1**). This specific expression pattern and the timing of polarization of both, GEF4 and GEF12, indicate similar functions during a defined time window in differentiating trichoblasts. This is distinctly different compared to GEF3 and GEF14, as these GEFs both polarize at a significantly earlier timepoint and show a more defined and restricted localization to the RHID.

**Figure 2.**
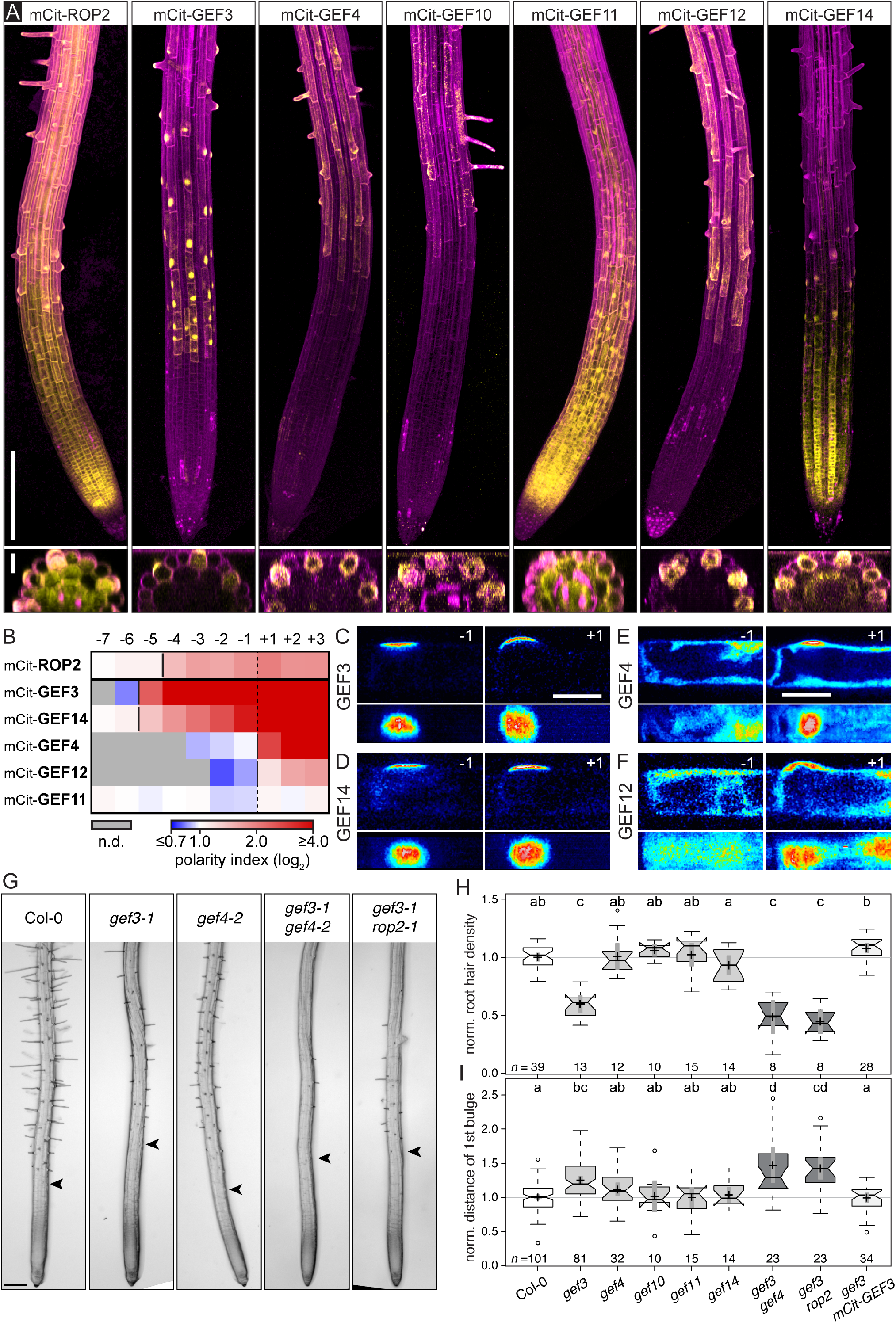
Functional analysis of RopGEFs in the root epidermis identifies GEF3 to be required for efficient root hair emergence. (**A**) Roots expressing mCit fusion proteins (yellow) under their respective promoters, stained with propidium iodide (magenta) to visualize root and cell outlines. Top panel: Maximum intensity projection of the root tip. Scale bar, 200 μm. lower Panel: Optical cross section through the root in the elongation zone to visualize the tissue-specific expression. Scale bar, 20 μm. (**B**) Timeline analysis of protein accumulation at the RHID for selected GEFs in comparison to ROP2. GEFs are sorted by timing of accumulation at the RHID. Polarity index measurement and calculation as in **Figure 1**. (**C-F**) Subcellular localization (intensity-coded representation) of mCit-GEF3, mCit-GEF4, mCit-GEF14 and mCit-GEF12, in one cell before (−1) and after bulging (+1), respectively. Confocal cross sections (top) and surface views (bottom) of the same cells are shown. Scale bars, 20μm. (**G**) Representative root tips of WT (Col-0), and *gef* single and double mutant lines. Arrowheads indicate the first bulge. (**H, I**) Quantification of root hair density (**H**) and distance (**I**) of first bulge to the root tip for selected single and double mutants. Values were normalized to corresponding Col-0 values for each independent experiment. Box-plots: number of measurements (n); center lines show medians; notches of boxes represent the 95% confidence interval of the medians; box limits indicate 25th and 75th percentiles as determined by R software; whiskers extend 1.5 times the interquartile range from the 25th and 75th percentiles, dots represent outliers; crosses represent sample means; grey bars indicate 95% confidence intervals of the means. Letters above box-plots show results of an ANOVA-Test (significance value *p*=0.01), with same letters indicating no significant differences. See also **Figure S2**.

In addition to the specific expression pattern of GEF3, its localization stood out in comparison to all other GEFs, as mCit-GEF3 appeared highly polarized at the cell periphery of the RHID, soon after the first fluorescence signal became detectable (**Figure 2A-C**). Polarization of mCit-GEF14 occurred less rapidly but reached a similar degree as mCit-GEF3 at stage +1 (**Figure 2B,D, Figure S1**), while mCit-GEF4 and mCit-GEF12, exhibited a lower degree of polarization and association with the plasma membrane (**Figure 2B, E, F, Figure S1**). mCit-GEF3 appeared highly concentrated in a circular polar domain, being barely detectable outside this domain (polarity index at cell “+1”: 14.4 ± 2.2) (**Figure 2C, Figure S1**). We tested whether the timing of GEF polarization was dependent on the timing of gene expression and analyzed the polarity indices of mCit-GEF3 and mCit-GEF4 under the control of estradiol-inducible promoters. In both cases, the earlier gene expression did not result in an earlier RHID polarization of the protein (**Figure S1**), but rather delayed the time point of significant polarization of mCit-GEF4, likely due to higher cytosolic levels that affected the polarity index calculation. Its independence from gene expression indicates that the timing of polarization is an intrinsic feature that may be controlled by further partners of unknown identity.

To determine the functional role of the investigated GEFs we analyzed T-DNA insertion lines for phenotypes in root hair initiation and development. Root hair density was used as a measure for the general ability to develop root hairs, distance of the first bulge from the root tip was used to infer delays in the timing of root hair initiation and root hair length was used as measure for regulation of tip growth. Only in *gef3* mutant alleles, root hair density was reduced and root hair initiation was delayed, (**Figure 2G-I**), while hair length seemed only slightly affected (**Figure S2E**). In *gef3-1*, which lacks almost all full length transcript (**Figure S2C**), the observed phenotypes could be rescued with a GEF3::mCit-GEF3 construct (**Figure 2H, I**), further verifying that loss of GEF3 function causes the phenotypes and confirming the functionality of the fusion construct. Furthermore, other mutant alleles of *gef3* showed a similar delay in root hair initiation (**Figure S2B,D**). None of the *gef3* T-DNA insertion lines lacked root hairs entirely, pointing towards functional redundancy among GEFs or compensatory mechanisms. A mutant line carrying both *gef3-1* and *rop2-1* alleles did not exhibit a significant increase in phenotype strength (**Figure 2G-I**), which indicates that such compensation involves other ROP-GEF combinations present in trichoblasts. Except for *gef3-1* no other *gef* single mutants, including *gef4-2*, exhibited any significant negative effects on bulge distance or root hair density (**Figure 2G-I**), but *gef4-2* lines did show reduced hair length (**Figure S2E**). A combination of *gef3-1* and *gef4-2* exhibited similar root hair density in *gef3-1*, but showed a small increase in distance of the first bulge from the root tip, i.e. stronger delay in hair initiation (**Figure 2G-I**). This phenotypic analysis suggests different, yet overlapping roles for GEF3 and GEF4, where GEF3 is mainly, but not exclusively, responsible for root hair initiation and bulging, while GEF4 is regulating root hair tip growth.

### GEF3 is required for efficient root hair emergence and polar recruitment of ROP2

We have shown that GEF3 accumulated very specifically at the RHID, preceded ROP2 accumulation, and was required for proper root hair development. Therefore, GEF3 was selected as a promising potential regulator of ROP dependent root hair initiation.

To determine the temporal relationship of GEF3 and ROP2 recruitment to the RHID in detail, we observed the localization of both proteins at higher resolution over time. We employed a new microfluidic device, termed RootChip-8S, for long-term imaging of growing *Arabidopsis* roots. The RootChip-8S, is an adapted version of previously published RootChip designs [51–53] and consists of eight separate observation chambers that can be individually perfused to provide stable growth conditions and restrict root movement in Z to prevent focal drift (**Figure S3A, B**). The first detectable signal of GEF3::mCit-GEF3 was observed in the transition zone of the root, shortly after the last cell division (**Figures 2 A and 3 A**). Time-lapse imaging showed early targeting of mCit-GEF3 to the rootward cell pole (**Figure 3 A**). As the protein gradually enriched at the RHID over approximately 30 min (**Figure 3 B**), its localization at the rootward cell pole vanished (**Figure 3 A**). The GEF3 domain eventually stabilized at a consistent diameter of 9.6 μm (± 2.5 μm; *n*=17; FWHM), with its center 9.0 μm (± 2.2 μm) from the rootward cell pole (**Figure 3 B, Figure S3C**). To validate the earlier RHID accumulation of GEF3 compared to ROP2 through simultaneous visualization in the same cells, we imaged their localization in growing roots co-expressing fluorescently labeled versions of both proteins and found that mCit-GEF3 accumulated at the RHID one stage earlier than mTurquoise2-ROP2 (mTq2-ROP2) (**Figure 3 C**).

**Figure 3.**
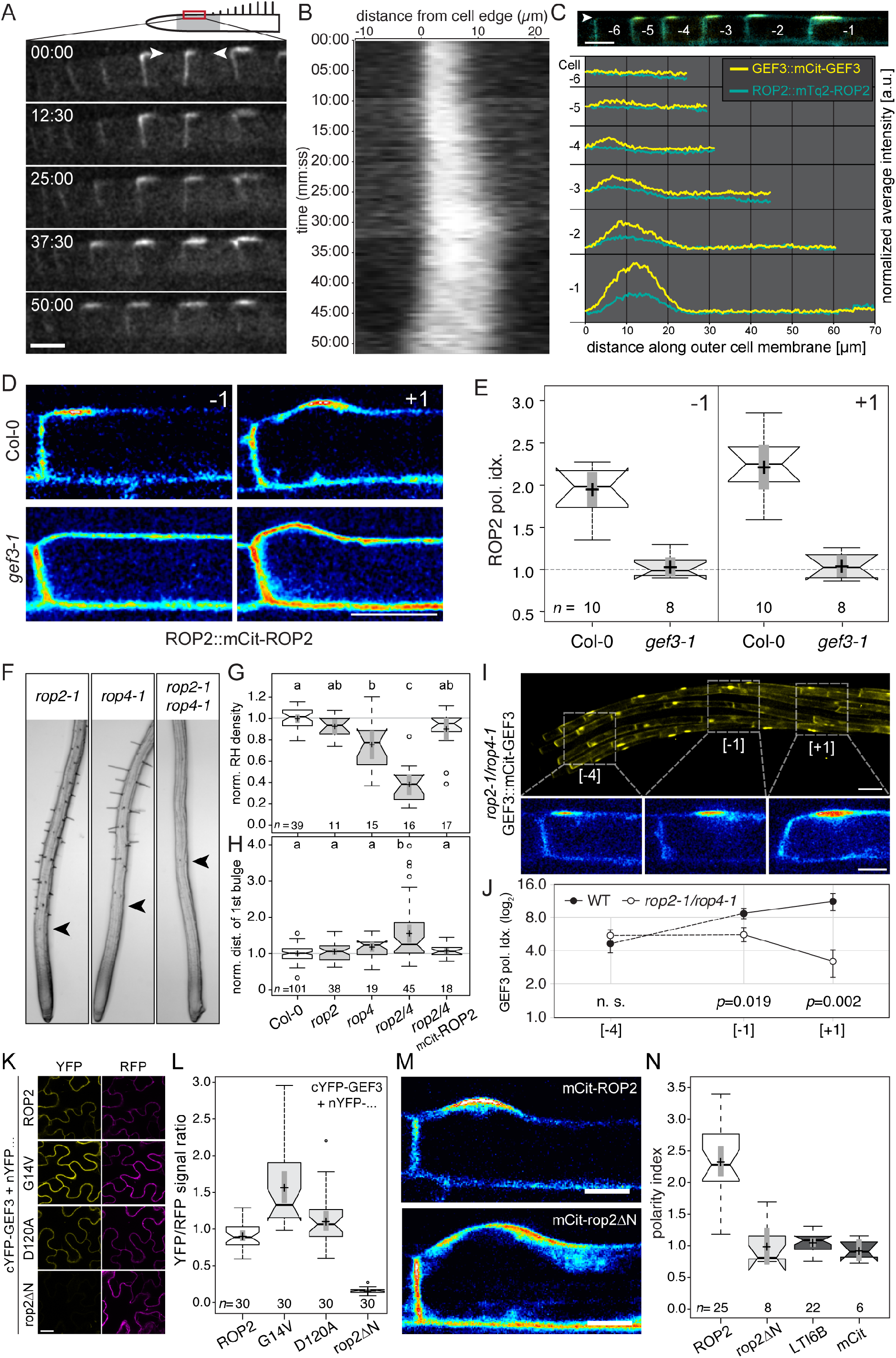
GEF3 is necessary for polar ROP2 recruitment during root hair initiation. (**A**) Time-lapse imaging of GEF3::mCit-GEF3 in the RootChip-8S (see **Figure S2**). A root hair cell file in the early elongation zone is shown. Multiple stages of protein accumulation can be seen in each still image. The time series was registered to compensate for cell movement due to root growth. Numbers indicate time (mm:ss). Scale bar, 20μm. (**B**) Kymograph of intensity distribution (horizontal axis) along the outer cell flank (left-right orientation) between arrow heads in (**A**) over time (vertical axis). Note the gradual expansion of the emerging mCit-GEF3 domain until approximately 30 min and a stable domain size thereafter. (**C**) Colocalization and intensity profiles of relative signal distribution for mCit-GEF3 and mTq2-ROP2 along root hair cell files. Top: example root hair cell file expressing ROP2::mTq2-ROP2 (cyan) and GEF3::mCit-GEF3 (yellow). Cells are labeled from youngest cells (−6) to oldest cell (−1) before bulging. Bottom: average profiles of relative mTq2-ROP2 (cyan) and mCit-GEF3 (yellow) intensities along the outer cell periphery (line width 3 px) of 6 root hair cell files (arrowhead indicates direction of line measurement). To compensate for intensity differences between measurements, the signal intensities of each construct are normalized to the value at position 0 μm of each cell and afterwards averaged. (**D**) Subcellular localization and (**E**) quantification of mCit-ROP2 polarity at the RHID in Col-0 and the *gef3-1* mutant in root hair cells one cell before (−1) and after bulging (+1). Scale bar, 20μm. (**F**) Representative root tips of *rop2-1, rop4-1* and *rop2-1/rop4-1* mutant lines. Arrowheads indicate the first bulge. (**G, H**) Quantification of root hair density and distance of first bulge to the root tip in ROP single and double mutants, as well as in *rop2-1/rop4-1* carrying a ROP2::mCit-ROP2 construct. Values are normalized to corresponding Col-0 values for each independent experiment (for comparison, data for Col-0 shown in Figure 2H,I are re-plotted). (**I**) GEF3::mCit-GEF3 expression and localization in *rop2-1/rop4-1* mutant background. Upper panel: overview of the elongation zone. Lower panel: subcellular localization of mCit-GEF3 in cells that correspond to stages −4, −1, and +1. Due to the lack of bulges in this mutant, the reference point used in other lines is missing here and stage designations are therefore estimates. (**J**) Quantification of the polarity index of GEF3::mCit-GEF3 in *rop2-1/rop4-1* mutant background. Stages are chosen as indicated in (**I**). (**K**) Representative images of an rBiFC assay in tobacco epidermal cells for each combination tested. Positive YFP signal indicates protein-protein interaction, RFP signal in the same cells serves as expression control (see Materials and Methods and **Figure S2** for details). All images were recorded using identical conditions and settings, correction for brightness and contrast during image processing was kept equal except for the exemplary YFP image of *rop2*ΔN, which was set brighter to visualize the very low background signal. (**L**) Quantification of ratiometric BiFC experiments. YFP signals were normalized to the corresponding RFP signal to allow comparison of interaction strength. (**M**) Confocal image of root hair cells expressing estradiol inducible mCit-*ROP2* or mCit-*rop2*Δ*N*. Scale bar, 10μm. (**N**) Quantification of polarity index of the constructs shown in (**N**) and compared to the non-polar membrane marker GFP-LTI6B and free mCitrine. For a detailed explanation of all shown box-plots, see the description given in the legend of Figure 2. See also **Figure S3**.

To test if GEF3 directly influences ROP2 recruitment to the RHID, we expressed mCit-ROP2 in the *gef3-1* T-DNA insertion mutant. Strikingly, we did not observe any significant polarization of mCit-ROP2 in the *gef3-1* mutant (**Figure 3 D, E**). Some cells in *gef3-1* were still able to produce bulges, but these bulges lacked polar ROP2 accumulation (**Figure 3 D, E**). Interestingly, the general membrane association of mCit-ROP2 was unaffected, as the localization appeared similar to that in other cell types. This shows that GEF3 function is required for ROP2 recruitment specifically to the RHID. We thus hypothesized that the observed root hair phenotypes in *gef3* mutants could be explained by failed polar recruitment and activation of downstream ROPs. Consequently, ROP loss-of-function mutants would exhibit similar phenotypes as *gef3*. We combined two previously described T-DNA lines for ROP2 and ROP4 [35, 54], and analyzed single and double mutants for phenotypes in root hair initiation. To account for the different genetic backgrounds of both single mutants (*rop2-1*, Col-0; *rop4-1*, WS), we transformed the double mutant *rop2-1/rop4-1* with a ROP2p::mTq2-ROP2 construct, and used the generated line as control for phenotype analysis. For both traits, root hair density and distance of the first bulge to the root tip, the double mutant *rop2-1/rop4-1* showed a phenotype that exceeded the strength of phenotypes in *gef3-1* (**Figure 3 F - H, Figure S2F,G**). The single mutant lines *rop2-1 and rop4-1* showed, however, no significant effects and our ROP2p::mTq2-ROP2 construct was able to fully revert phenotypes in *rop2-1/rop4-1*, demonstrating functional redundancy among both ROPs. In contrast to the loss of ROP2 polarity in *gef3-1* mutants, polar mCit-GEF3 domains were still observed in *rop2-1/rop4-1* mutants (**Figure 3 I, J**). mCit-GEF3 polarity was unaffected in the early elongation zone (**Figure 3 I, J; stage −4**), but was slightly reduced in later stages (**Figure 3 I, J; stages −1, +1**) compared to mCit-GEF3 polarity in Col-0 background (**Figure S1**). This indicates that initial GEF3 polarization is largely ROP independent, while ROPs have a positive effect on GEF3 polarity in later stages of root hair development. The question remained how GEF3 fulfills its function as a factor directing polarization of ROP2. Previous studies have demonstrated that GEF-catalyzed nucleotide exchange of ROPs involves the formation of a stable G protein-GEF complex [55–58]. To test if GEF3 was capable of recruiting ROP2 through physical interaction, we used a ratiometric bimolecular fluorescence complementation (rBiFC) assay with GEF3 fused to a C-terminal fragment of YFP (cYFP) and either an N-terminal fragment of YFP (nYFP)-carrying ROP2 (wild-type), a constitutively active CA-ROP2 (G14V), a dominant-negative DN-ROP2 (D120A), or a truncated *rop2*ΔN (Δ1-79) that lacks its interaction domain (**Figure S3D**) [5, 59], co-expressed transiently in *Nicotiana benthamiana* leaf epidermis cells [60]. All GEF3-cYFP/nYFP-ROP2 combinations, except for *rop2*ΔN, resulted in detectable YFP fluorescence (**Figure 3 K, L, Figure S3H-J**). This finding was corroborated by a split-ubiquitin assay in yeast [61], testing the same putative interactor combinations. Except for *rop2*ΔN, all forms of ROP2 yielded yeast growth, indicating physical interaction with GEF3 (**Figure S3E-G**). Consistent with the hypothesis that physical interaction with GEF3 is required for polar recruitment of ROP2 to the RHID, mCit-*rop2*ΔN expressed in *Arabidopsis* trichoblasts did not assume polar localization at the RHID (stage +1, **Figure 3 M, N**). Our interaction assays show further that GEF3 physically interacts with the active as well as the inactive form of ROP2 via its interaction domain. While physical interaction between GEF3 and CA-ROP2 was unexpected at first, specific binding of certain RopGEFs to both, GDP- and GTP-bound ROPs has been demonstrated before [43].

Taken together, these results show that GEF3 is necessary for ROP polarization by recruiting ROP2 to the RHID via direct physical interaction. Furthermore, the presence of ROP2 and ROP4 is not needed for GEF3 polarization, but enhances GEF3 polarity.

### GEF3 defines the RHID and guides ROP polarization

To investigate whether GEF3 is not only necessary but also sufficient for ROP2 polarization, we studied transgenic lines, inducibly overexpressing mCit-GEF3 (mCit-GEF3ox) in Col-0 or in ROP2::mTq2-ROP2 background. Upon induction of mCit-GEF3 expression we observed that additional patch-like GEF3 domains formed in trichoblasts and, surprisingly, also in atrichoblasts (**Figure 4 A**). We frequently found mCit-GEF3 accumulation at the shootward sides of cells in the meristem or the early elongation zone (**Figure 4 A**). Importantly, all observed normal and ectopic mCit-GEF3 accumulations resulted in the simultaneous recruitment of mTq2-ROP2 (**Figure 4 A**).

**Figure 4.**
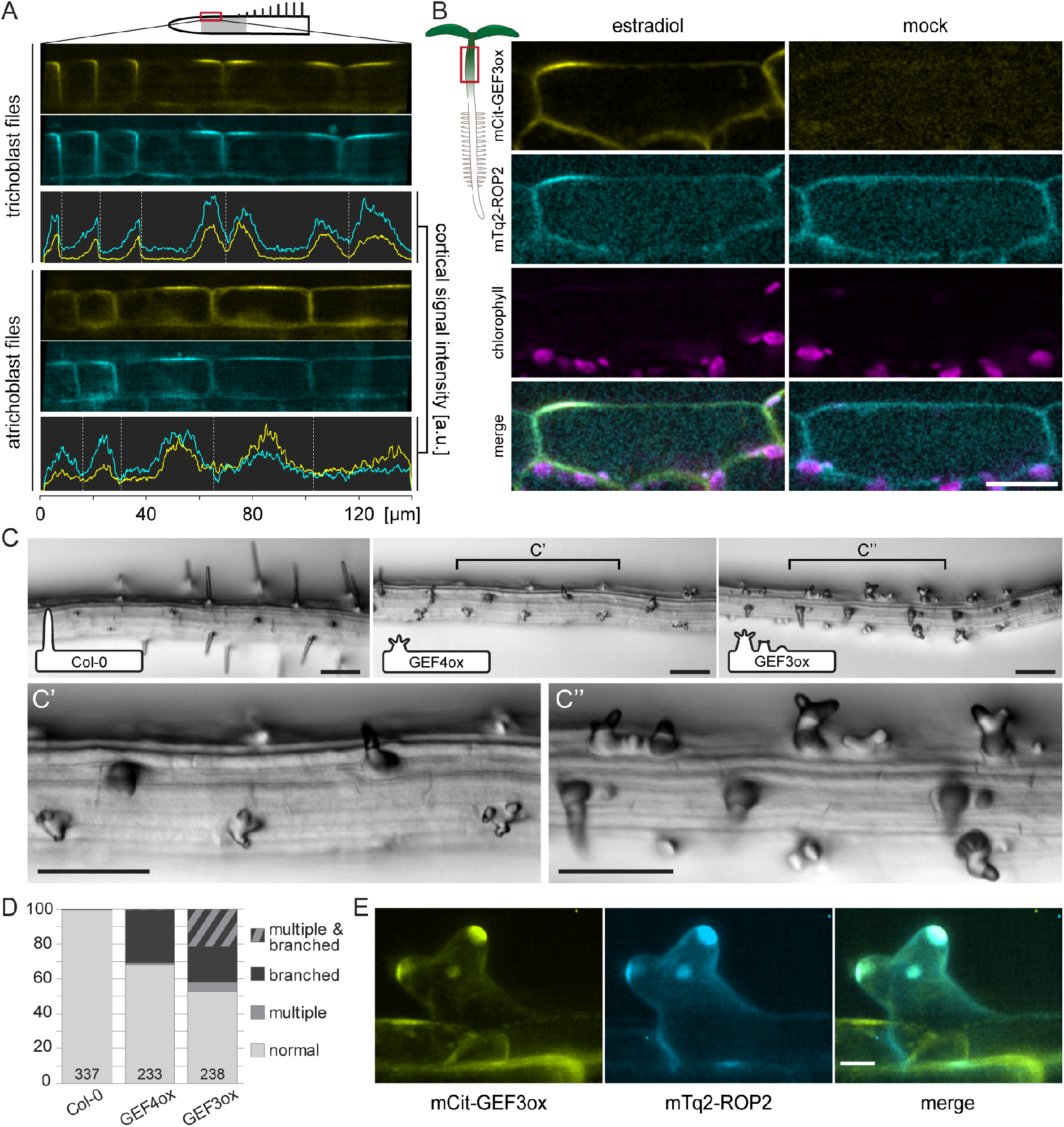
GEF3 initiates the formation of polar growth domains. (**A**) Localization of *ROP2*::mTq2-ROP2 (cyan) in roots overexpressing mCit-GEF3 (yellow). Top: representative cell files of trichoblasts and atrichoblasts, both show ectopic accumulations of both markers. Bottom: intensity profiles along cell periphery of cells shown on the left. Dashed lines indicate cell borders. (**B**) Subcellular localization of *ROP2*::mTq2-ROP2 (cyan) in hypocotyl cells ectopically expressing mCit-GEF3 (yellow). To visualize the background signal in the mCit channel of the mock-treated seedling, intensities were 10-fold enhanced during image processing as compared to the image of estradiol-treated specimen. Other channels are scaled identically. Scale bar, 20 μm. (**C**) Top panels: Representative images of Col-0 (left) and roots overexpressing mCit-*GEF4* (middle) and mCit-*GEF3* (right) (24h after estradiol induction). Inserts show an idealized sketch of the predominant root hair shape. Bottom panels: Higher magnification views of the root zones indicated as C’ and C” in the top panels of GEF4ox and GEF3ox, respectively. Scale bars, 100 μm. (**D**) Frequency of phenotypes (induction of multiple hairs, branched bulges, and combined phenotypes) induced by overexpression of lines shown in (**C**) in % of total observed hairs. Bottom numbers indicate *n*. (**E**) Colocalization of mCit-GEF3 (yellow) and ROP2::mTq2-ROP2 (cyan) in branched root hairs, 24 hours after estradiol induction of mCit-GEF3 overexpression. Scale bar, 20 μm. See also **Figure S4**.

We took advantage of the ubiquitous overexpression of mCit-GEF3 and determined its localization in cells outside the root. In epidermal cells of the hypocotyl, we occasionally observed occurrences of ectopic GEF3 accumulation into domains that strikingly resembled those of RHIDs in trichoblasts (**Figure 4 B, Figure S4A**). Consistently, these ectopic mCit-GEF3 domains also lead to recruitment of mTq2-ROP2, which never occurred in hypocotyl cells under control conditions (**Figure 4 B**). Except for trichoblasts, mCit-GEF3ox did not lead to bulge formation in other cells, neither did induced expression of mCit-GEF3ox influence the timing of bulging (**Figure S4B**), indicating that ROP polarization alone was not sufficient to trigger outgrowth and further trichoblast-specific factors are required.

Since the analysis of gene expression, protein localization and loss-of-function phenotypes all suggested a differential timing of GEF3 and GEF4 function, we hypothesized that gain-of-function phenotypes should also reflect their specific roles in root hair initiation and growth. We therefore compared the phenotypic effects of mCit-GEF3ox to mCit-GEF4ox, which is known to induce root hair branching in trichoblasts [62]. In mCit-GEF4ox lines, bulges with branched tips, which failed to form long tube-like hairs, occurred in 30.5% of the observed cells (**Figure 4 C, D**). mCit-GEF3ox roots also displayed root hairs with branched tips in 20.6% of all trichoblasts, resembling mCit-GEF4 overexpression (**Figure 4 C, D**). However, mCit-GEF3ox further led to the formation of additional bulges, with 5.5% of trichoblasts showing multiple unbranched bulges and 21.4% of the cells showing multiple branched bulges within one cell (**Figure 4 C, D**). This phenotype bears striking similarities to *scn1* mutants that lack a ROP-inhibiting RhoGDI [11], which further points towards a gain of ROP activity in mCit-GEF3ox roots at an early stage during RHID positioning. We observed mCit-GEF3 accumulations in tips of branched root hairs and mTq2-ROP2 was generally present in the same locations (**Figure 4 E**), thus confirming that mCit-GEF3ox was also responsible for the formation of additional polar domains in branching hairs. These differential effects of GEF3 and GEF4 overexpression resemble the differential timing of their polar localization and further support the hypothesis that both proteins serve specific functions during different phases of root hair development, with GEF3 being decisive for the early formation of the RHID and ROP2 recruitment, and GEF4 being involved in downstream activation of ROP2 to trigger growth.

### GEFs serve as positional cues during cell polarization

To our knowledge, GEF3 is not only the first RopGEF member reported to temporally precede ROP polarization during tip growth initiation, but it is also the first example for a plant protein that, when overexpressed, is capable of inducing the ectopic formation of polar, ROP-recruiting domains. Our finding, that GEF3 acts upstream of ROP2, challenges the idea of ROP self-organization during the initiation of cell polarization in trichoblasts [63]. A self-organizing reaction-diffusion mechanism, involving the interplay of ROP-activating RopGEFs and ROP-inactivating RopGAPs has been found to underlie the regular cell wall patterning in differentiating metaxylem vessels [64, 65]. In contrast to the multiple, regularly spaced cell wall pits of the metaxylem, the dependable positioning and establishment of a single polar RHID in trichoblast is likely to require a mechanism that eliminates randomness during its formation. Our results suggest that such non-random polarization can be achieved through a GEF3-mediated recruitment and temporally controlled ROP activation by differentially timed GEFs at a predefined location.

Evidence for a GTPase-positioning function of GEFs can also be found in other organisms. Recent work shows that during cellularization in Drosophila embryos, the GEF Dizzy and the heteromeric GEF complex ELMO-Sponge are required for polarization of the GTPase Rap1 [41, 66]. It remains, however, challenging to establish the molecular sequence of GEF and GTPase functions in the context of a developing embryo. In budding yeast, Cdc42 is activated by its GEF Cdc24 during both budding and shmoo formation [67–69]. To initiate budding in yeast, the GEF Bud5 is recruited to the edge of the previous cell-division site by the landmark Axl2 [70]. Stabilized by scaffold proteins, Bud5 and the GEF Cdc24 then locally activate the GTPases Rsr4 and Cdc42 [71–76].

Using a GEF-scaffold complex as landmark for the targeted deployment of the growth machinery may therefore represent a common design principle among eukaryotic cells to prevent spontaneous and random polarization and, instead, drive growth in a directed manner. Yet, the fact that RopGEFs share no conserved features with animal or fungal GEFs [42], indicates that the problem of robust cell morphogenesis has been solved several times during evolution to account for specific differences in cellular growth mechanisms, organ development and lifestyle.

## ACKNOWLEDGMENTS

We are grateful to Dominique Bergmann (Stanford) and Sheila McCormick (UC Berkeley) for detailed comments on our manuscript. We thank Karin Schumacher (U. Heidelberg) and Thomas Dresselhaus (U. Regensburg) for helpful advice, all members of the Grossmann lab and our colleagues at COS Heidelberg for inspiring discussions and their active support. We thank Jan Lohmann (Heidelberg), Markus Grebe (Potsdam), Masa Sato (Kyoto), Tijs Ketelaar (Wageningen), Kateřina Schwarzerová (Prague), Yvon Jaillais (Lyon), and Ueli Grossniklaus (Zurich) for sharing materials and Karin Schumacher and Thomas Holstein (both Heidelberg) for access to microscopes. This work was supported by research group funds from the excellence cluster CellNetworks to G.G., grants from the Deutsche Forschungsgemeinschaft (GR 4559/3-1 to G.G. and EV 198/1-1 to J.F.E.), seed funding through the SFB1101, an Emmy Noether fellowship (GR 4251/1-1) to C.G., a PhD-student grant from the Carl-Zeiss-Stiftung to D.G.M., and financial support by ETH Zürich to C.E.S.

## AUTHOR CONTRIBUTIONS

The study was conceived by P.D. and G.G.; P.D., A.R., V.A.F.S., D.G.M., L.Y.A., and N.F.K. performed experiments; P.D., A.R., V.A.F.S., D.G.M., L.Y.A., C.G. and G.G. analyzed data. C.E.S. and G.G. designed the microfluidic device. J.F.E. designed the spinning disk microscope and gave advice about the imaging procedure. P.D., A.R. and G.G. prepared figures and wrote the manuscript with input from all coauthors. All authors read and approved the final version of the manuscript.

## MATERIALS AND METHODS

### Plant material

*Arabidopsis thaliana* ecotype Col-0 was used as wildtype in this study. The mutant lines *rop2-1* (SALK_055328C), *gef3-1* (SALK_079879C), *gef3-2* (SALK_079885), *gef3-3* (SAIL_294_A09), *gef3-4* (SALK_021751), *gef3-5* (SALK_046978C), *gef4-2* (SALK_107520) *gef10-1* (SALK_009456), *gef11-1* (SALK_126725C), *gef14-2* (SALK_046067) were obtained from NASC (Nottingham Arabidopsis Stock Centre http://arabidopsis.info/). The *rop4-1* mutant [36] was kindly provided by Markus Grebe (Potsdam). Double mutants were made by crossing and selection for double homozygous plants by PCR (Primers used for genotyping, see Table S3). The GFP-SYP123 and GFP-SYP132 lines [26] were kindly provided by Masa H. Sato (Kyoto), SEC3A-GFP lines [27] were kindly provided by Tijs Ketelaar (Wageningen), the GFP-ARPC2A line [25] was kindly provided by Kateřina Schwarzerová (Prague), and the PI(4,5)P_2_-reporter line (P15Y) [31] was kindly provided by Yvon Jaillais (Lyon). To generate transgenic Col-0 lines expressing GFP-tagged FER, Col-0 was transformed with pFER::FER-GFP containing pMDC111 [77], kindly provided by Ueli Grossniklaus (Zurich).

Information on all genes referenced in this work, including mutant alleles and sources is provided in **Supplementary Data S1, Table S1**.

### RNA extraction and expression analysis

Seedlings were grown on mesh on a ¼ MS plate. Root tissue was harvested 10 dag, immediately frozen in liquid nitrogen and ground in a tissue lyser (Qiagen) with metal beads. Total RNA was extracted using the RNeasy Plant Mini Kit (Qiagen). After DNAse digest (Thermo Fischer Scientific), 0.5 μg total RNA was used for cDNA synthesis with oligo(dT) primers (RevertAid First Strand cDNA Synthesis Kit, Thermo Fischer Schientific). To test for the presence of Gef3 mRNA, 2μl of this cDNA was used for PCR amplification (40 cycles). The primers used for the expression analysis are listed in Supplementary Information, **Table S2**.

Data mining of published cell type-specific gene expression was performed using Genevestigator [78].

### Growth conditions and inducible expression

Plants were grown under long day conditions (16 h light) at 21°C. Seedlings used for experiments were grown for seven days on ¼ MS medium (pH 5.7) without sucrose, supplied with 0.8% plant agar (Duchefa). Transgenic plants transformed with an estradiol inducible construct were induced by covering with cellulose tissues soaked with ¼ MS medium containing 20 μM Estradiol (Stock 20 mM in Ethanol) and 0.01% Silwet L77 for 15 minutes. The time after induction differed between the individual expression constructs and ranged from 3 – 12 h to get sufficient signal for imaging. In general, for imaging of protein localization using inducible expression constructs, time after induction was as low as possible to avoid overexpression artefacts.

### Plant transformation and selection

The agrobacterium strain GV3101::pMP90RK pre-transformed with the helper plasmid pSoup was used for all transformations of *Arabidopsis thaliana*. Plants were transformed using floral dipping and transgenic T1 plants were recovered after the appropriate selection. Selection of constructs containing mTurquoise2 (mTq2) was done on plates containing 50 μg/ml Kanamycin. T1 Selection of constructs containing mCitrine (mCit) was done on soil spraying 3x within 2 weeks (7 – 21 day old seedlings) with Basta^®^-Solution containing 200 μg/ml Glufosinate-ammonium and 0.05% Tween20. Further generations were selected on plates containing Basta^®^-Solution to a concentration of 7.5 μg/ml Glufosinate-ammonium. At least 10 independent transgenic lines per expression construct were recovered and checked for fluorescence. Lines that had clear fluorescence and limited phenotypic anomalies were confirmed by PCR and used for further study.

### Molecular cloning

All cloning for expression vectors used in Arabidopsis was performed using the GreenGate system [79] and base vector pGGZ003 was used for all expression constructs. A List of all expression constructs including module composition and tagging site is provided in Supplementary Information, **Table S3**. Where internal BsaI (Eco31I) sites were present, they were mutated fusing two PCR fragments, with one mutating the internal BsaI site by an altered primer sequence at this site. After cloning the individual modules into the entry vectors, the complete fragment was checked by sequencing. To confirm the correct assembly of modules into pGGZ003, the presence and orientation was checked by sequencing. As fluorophores, mCitrine (mCit) [80] and mTurqoise2 (mTq2) [81] were fused to the gene of interest separated by a short GAGAGA-Linker. mCit was cloned from the Ca^2+^ indicator CerTN-L15, containing the common folding mutations V163A and S175G [82], and made monomeric by introducing the A206K mutation. mTq2 was cloned from the Addgene vector pLifeAct-mTurquoise2 (ID #36201). A List of all primers used for molecular cloning is provided in Supplementary Information, **Table S2**.

For estradiol inducible expression constructs [83], the XVE gene was fused to the promoter (635 bp) and terminator of *UBI10* (AT4G05320). This expression cassette was cloned in reverse orientation into the entry vector pGGA000 together with the oLexA minimal 35S promoter element. Both elements were separated by a 560 bp spacer. This resulted in a single module that could be used as a promoter module for GreenGate cloning.

For expression of genes under control of their native promoter, the following sequences upstream of the start codon were cloned into pGGA000: *GEF3* −1743 bp; *GEF4* −1500 bp; *GEF10* −1534 bp; *GEF11* −1458 bp; *GEF12* −701 bp; *ROP2* −1503 bp; *PIP5K3* −1193 bp.

For the mbSUS and rBiFC analysis, constructs (listed in Supplementary Information, **Table S3**) were generated using Gateway technology or the 2in1 cloning system (Grefen and Blatt 2012). Coding sequences were PCR-amplified and inserted via ‘BP’ recombination into either pDONR207, pDONR221-P1P4, or pDONR221-P3P2, respectively, and confirmed by sequencing. Resulting ‘Entry constructs’ were cloned via ‘LR’ recombination into Destination vectors for either mbSUS or rBiFC [84].

### Phenotyping of T-DNA insertion lines and inducible overexpression mutant lines

Primary roots (7 dag) were used for all phenotyping experiments. For phenotyping of T-DNA insertion lines the distance of the first bulge to the root tip was defined. The first visible swelling of the cell outline was defined as first bulge, even when this morphological change was very little. The root hair density was analyzed in the next 2 mm. The root hair length was analyzed measuring root hairs growing perpendicular to the root in a region 3-6 mm away from the root tip. For phenotyping the overexpression lines, roots were induced by spraying with medium containing 20 μM Estradiol. The position of the root tip was marked just after induction and normal growing roots were imaged and analyzed 24-26 hours after induction (h. a. i.). A region approximately 2-4 mm from the root tip was used for the analysis of the overexpression phenotype.

### Propidium iodide (PI) staining

Primary roots (7 dag) were stained for 5 min in 10 μg/ml propidium iodide (PI; diluted in ¼ MS, from 1 mg/ml stock), washed in medium and imaged directly. As PI staining varies substantially between different developmental zones of the root, we employed gamma correction (0.75) exclusively in the PI channel of the images to equalize stronger and weaker stained regions in the final figures (see image processing).

### Ratiometric Bimolecular Fluorescence analyses (rBiFC)

Agrobacterium-mediated transient transformation of 4-6 week old *N. benthamiana* leaves with 2in1 rBiFC constructs was performed as described previously [85]. Fluorescence intensities were recorded 36h post infiltration for approximately 30 images per construct as described in [60]. Ratios between complemented YFP fluorescence and RFP were calculated and plotted using BoxplotR. Immunoblotting verified protein expression utilizing anti-MYC conjugated to peroxidase antibodies and anti-HA peroxidase conjugated antibodies, respectively.

### Mating-based Split Ubiquitin Assay (‘Cyto-SUS’)

Bait/Cub fusion and preys/Nub fusions were generated by cloning the coding region for *GEF3* and different *ROP2* versions into the Gateway-compatible vectors pMetOYC-Dest and pNX35-Dest, respectively. These were transformed into haploid yeast strains THY.AP4 (Cub fusion) and THY.AP5 (Nub fusions) as described previously [86]. After mating, diploid yeasts were dropped in 10 times OD dilutions on vector- (CSM-Leu-, Trp-, Ura) and interaction-selective media (CSM-Leu-, Trp-, Ura-, Ade-, His-) supplemented with 50 or 500 μM methionine. The non-mutated N-terminal ubiquitin moiety (NubI) was used as positive control, and NubI13G (empty vector) as negative control [84]. Protein expression was verified in haploid yeast using antibodies against the VP16 domain for the Cub fusion or the HA epitope tag for the Nub fusions.

### Microfluidic device (RootChip-8S) fabrication and on-chip plant cultivation

Microfluidic device fabrication and on-chip plant cultivation were performed as described in [87, 88]. Briefly, Arabidopsis seedlings were grown in pipette tips filled with solidified medium and interfaced with poly(dimethylsiloxane) (PDMS) RootChip-8S devices 4-5 days after germination under sterile conditions. Media perfusion through the microfluidic channels was controlled using a peristaltic pump (DNE GmbH; volumetric flow rate in each channel, 5 μl/min).

### Bright-field and fluorescence imaging

Live-cell fluorescence imaging was performed on a custom-built spinning-disk confocal microscope, consisting of a Nikon Ti-E stand, equipped with 20x multi-immersion (N.A. 0.75) and 60x water immersion (N.A. 1.2) objectives (Nikon), a motorized stage (Applied Scientific Instrumentation, USA), a spinning disk with customized hole pattern (CREST Optics, Italy), a motorized filter wheel (Cairn Research, UK), a laser launch (Omicron, Germany), and an EMCCD camera (Photometrics, USA). Image acquisition was operated through Nikon NIS Elements software or Micro-Manager [89]. mCitrine-labeled proteins were excited using a laser emitting at the wavelength of 515 nm; mTq2-labeled proteins were excited at 440 nm. Bandpass filter specifications were CWL (central wavelength) = 542 nm, BW (band width) = 20 nm for mCit emission and CWL = 480 nm, BW = 40 for mTq2. Chlorophyll autofluorescence was recorded using 561 nm excitation and emission filter specifications of CWL = 630 nm, BW = 92. Root hair phenotyping was performed by transmitted light microscopy on a Nikon SMZ18 stereo microscope, equipped with SHR Plan Apo 0.5x (N.A. 0.075) and 2x (N.A. 0.3) objectives (Nikon) and an Orca Flash 4.0 sCMOS camera (Hamamatsu, Japan). Imaging of the propidium iodide stained root tips expressing different *GEF*::mCit-*GEF* fusions was performed on a Leica SP5 point scanning confocal microscope with a 20x water immersion objective (N.A 0.7), using 1x Zoom, Pinhole 1AU, 1024×1024 pixel scanning field, 400 Hz scanning speed, 2x line average as settings. mCit was excited with the 514 nm line of an Argon-Laser and emission light was collected between 520-550 nm. Propidium iodide was exited with a 561 nm laser and emission light was collected between 570-650 nm. Both channels were imaged in sequential scans to avoid crosstalk using Hybrid-Detectors (Gain 200). Z-Stacks were acquired with 2 μm step size for 20 X objectives and 0.6 μm for 60/63 X objectives. The top-view of the RHID was reconstructed from Z-Stacks using the built-in function of FIJI (3D-Project with interpolation).

### Image processing, data analysis and software

All image processing, image analysis and measurements was done using FIJI (ImageJ) [90], further data analysis was done in Microsoft Excel, if not mentioned otherwise. A gaussian filter (radius 0.8 px) and a rolling ball (radius 100 px) background subtraction was applied to all shown images. The “Royal” lookup table was used for images shown in intensity pseudo color. Kymographs were generated using the multi-kymograph plugin in FIJI. For images of propidium iodide (PI) staining for visualization of the root outlines, the gamma settings were adjusted to 0.75 to reduce the contrast between weakly and strongly stained regions of the root. The adjustment was only done in the PI channel. The signal of the mCitrine-fusions was not altered.

To obtain vertical root cross sections, Z-stacks (voxel size: 0.7568×0.7568×2.0142 μm^3^) were processed using the Reslice function in FIJI. Maximum intensity projections of approximately 100 μm along the root axis, in a distance of approximately 700 to 1000 μm away from the root tip were generated.

Polarity indices were analyzed on the original unprocessed images by measuring in 3 by 15 pixel regions at the RHID, outside the RHID and outside the root, as a background value. Before bulge formation a region approximately 10 μm from the rootward cell edge was used as the RHID. The background value was subtracted from both regions and the value outside the RHID was divided by the value inside the RHID [(Int_in_ - Int_bkgd_) / Int_out_ - Int_Bkgd_)]. This polarity index was calculated for each cell within a root hair file, all files were aligned to the first cell with a visible bulge and the average polarity index for each developmental step was calculated. A signal was defined as polar, if its value differed (p-value <0.05, according to two-tailed t-test with unequal variances) to the polarity index of the membrane marker GFP-LTI6B.

To measure GEF3 domain size, intensity profiles along the outer plasma membrane were generated from confocal cross sections of trichoblasts. Since gradually decreasing fluorescence signals impede an unambiguous determination of the outer boundary of polar accumulations, domain size was quantified by measuring the <u>f</u>ull signal <u>w</u>idth at <u>h</u>alf-<u>m</u>aximal signal intensity (FWHM; **Figure S3**).

Box plots were created using BoxPlotR (http://shiny.chemgrid.org/boxplotr/), an R-based web-tool provided by the Tyers (Montreal) and Rappsilber (Edinburgh) labs. For all plots number of measurements (n) is given. Center lines show the medians; notches of boxes represent the 95% confidence interval of the medians; box limits indicate the 25th and 75th percentiles as determined by R software; whiskers extend 1.5 times the interquartile range from the 25th and 75th percentiles, outliers are represented by dots; crosses represent sample means; bars indicate 95% confidence intervals of the means. As statistical test, t-tests (two-tailed, unequal variances) were performed in Microsoft Excel, ANOVA tests were performed in R and the used script is provided in **Data S1**.

## SUPPLEMENTAL INFORMATION

**Table S1**. List of identifiers for genes and single mutant alleles used in this study.

**Table S2**. List of PCR primers used for GreenGate and Gateway cloning.

**Table S3**. List of used expression vectors.

**Data S1**. R script for ANOVA/Tukey tests performed on mutant phenotype data.

## REFERENCES

1. Riquelme, M. (2013). Tip growth in filamentous fungi: a road trip to the apex. Annu Rev Microbiol 67, 587–609.

2. Rounds, C. M., and Bezanilla, M. (2013). Growth Mechanisms in Tip-Growing Plant Cells. Annu Rev Plant Biol.

3. Russell, S. A., and Bashaw, G. J. (2017). Axon guidance pathways and the control of gene expression.

4. Molendijk, A. J., Bischoff, F., Rajendrakumar, C. S., Friml, J., Braun, M., Gilroy, S., and Palme, K. (2001). Arabidopsis thaliana Rop GTPases are localized to tips of root hairs and control polar growth. EMBO J 20, 2779–2788.

5. Jones, M. A., Shen, J.-J., Fu, Y., Li, H., Yang, Z., and Grierson, C. S. (2002). The Arabidopsis Rop2 GTPase is a positive regulator of both root hair initiation and tip growth. Plant Cell 14, 763–776.

6. Bi, E., and Park, H.-O. (2012). Cell polarization and cytokinesis in budding yeast. Genetics 191, 347–387.

7. Ikeda, Y., Men, S., Fischer, U., Stepanova, A. N., Alonso, J. M., Ljung, K., and Grebe, M. (2009). Local auxin biosynthesis modulates gradient-directed planar polarity in Arabidopsis. Nat Cell Biol 11, 731–738.

8. Fischer, U., Ikeda, Y., Ljung, K., Serralbo, O., Singh, M., Heidstra, R., Palme, K., Scheres, B., and Grebe, M. (2006). Vectorial information for Arabidopsis planar polarity is mediated by combined AUX1, EIN2, and GNOM activity. Curr Biol 16, 2143–2149.

9. Kiefer, C. S., Claes, A. R., Nzayisenga, J.-C., Pietra, S., Stanislas, T., Hüser, A., Ikeda, Y., and Grebe, M. (2015). Arabidopsis AIP1-2 restricted by WER-mediated patterning modulates planar polarity. Development 142, 151–161.

10. Fu, Y., Li, H., and Yang, Z. (2002). The ROP2 GTPase controls the formation of cortical fine F-actin and the early phase of directional cell expansion during Arabidopsis organogenesis. Plant Cell 14, 777–794.

11. Carol, R. J., Takeda, S., Linstead, P., Durrant, M. C., Kakesova, H., Derbyshire, P., Drea, S., Zárský, V., and Dolan, L. (2005). A RhoGDP dissociation inhibitor spatially regulates growth in root hair cells. Nature 438, 1013–1016.

12. Chang, F., Gu, Y., Ma, H., and Yang, Z. (2013). AtPRK2 Promotes ROP1 Activation via RopGEFs in the Control of Polarized Pollen Tube Growth. Mol Plant 6, 1187–1201.

13. Hwang, J.-U., Gu, Y., Lee, Y.-J., and Yang, Z. (2005). Oscillatory ROP GTPase activation leads the oscillatory polarized growth of pollen tubes. Mol Biol Cell 16, 5385–5399.

14. Hwang, J.-U., Vernoud, V., Szumlanski, A., Nielsen, E., and Yang, Z. (2008). A tip-localized RhoGAP controls cell polarity by globally inhibiting Rho GTPase at the cell apex. Curr Biol 18, 1907–1916.

15. Lin, Y., Wang, Y., Zhu, J. K., and Yang, Z. (1996). Localization of a Rho GTPase Implies a Role in Tip Growth and Movement of the Generative Cell in Pollen Tubes. Plant Cell 8, 293–303.

16. Yalovsky, S., Bloch, D., Sorek, N., and Kost, B. (2008). Regulation of Membrane Trafficking, Cytoskeleton Dynamics, and Cell Polarity by ROP/RAC GTPases. Plant Physiol 147, 1527–1543.

17. Zhang, Y., and McCormick, S. (2007). A distinct mechanism regulating a pollen-specific guanine nucleotide exchange factor for the small GTPase Rop in Arabidopsis thaliana. Proc Natl Acad Sci USA 104, 18830–18835.

18. Kost, B., Lemichez, E., Spielhofer, P., Hong, Y., Tolias, K., Carpenter, C., and Chua, N. H. (1999). Rac homologues and compartmentalized phosphatidylinositol 4, 5-bisphosphate act in a common pathway to regulate polar pollen tube growth. J Cell Biol 145, 317–330.

19. Cutler, S. R., Ehrhardt, D. W., Griffitts, J. S., and Somerville, C. R. (2000). Random GFP::cDNA fusions enable visualization of subcellular structures in cells of Arabidopsis at a high frequency. Proc Natl Acad Sci USA 97, 3718–3723.

20. Kusano, H., Testerink, C., Vermeer, J. E. M., Tsuge, T., Shimada, H., Oka, A., Munnik, T., and Aoyama, T. (2008). The Arabidopsis Phosphatidylinositol Phosphate 5-Kinase PIP5K3 is a key regulator of root hair tip growth. Plant Cell 20, 367–380.

21. Foreman, J., Demidchik, V., Bothwell, J. H., Mylona, P., Miedema, H., Torres, M. A., Linstead, P., Costa, S., Brownlee, C., Jones, J. D., et al. (2003). Reactive oxygen species produced by NADPH oxidase regulate plant cell growth. Nature 422, 442–446.

22. Takeda, S., Gapper, C., Kaya, H., Bell, E., Kuchitsu, K., and Dolan, L. (2008). Local positive feedback regulation determines cell shape in root hair cells. Science 319, 1241–1244.

23. Wong, H. L., Pinontoan, R., Hayashi, K., Tabata, R., Yaeno, T., Hasegawa, K., Kojima, C., Yoshioka, H., Iba, K., Kawasaki, T., et al. (2007). Regulation of Rice NADPH Oxidase by Binding of Rac GTPase to Its N-Terminal Extension. Plant Cell 19, 4022–4034.

24. Mendrinna, A., and Persson, S. (2015). Root hair growth: it’s a one way street. F1000Prime Rep 7, 23.

25. Havelková, L., Nanda, G., Martinek, J., Bellinvia, E., Sikorová, L., Šlajcherová, K., Seifertová, D., Fischer, L., Fišerová, J., Petrásek, J., et al. (2015). Arp2/3 complex subunit ARPC2 binds to microtubules. Plant Science 241, 96–108.

26. Ichikawa, M., Hirano, T., Enami, K., Fuselier, T., Kato, N., Kwon, C., Voigt, B., Schulze-Lefert, P., Baluska, F., and Sato, M. H. (2014). Syntaxin of Plant Proteins SYP123 and SYP132 Mediate Root Hair Tip Growth in Arabidopsis thaliana. Plant Cell Physiol 55, 790–800.

27. Zhang, Y., Immink, R., Liu, C.-M., Emons, A.-M., and Ketelaar, T. (2013). The Arabidopsis exocyst subunit SEC3A is essential for embryo development and accumulates in transient puncta at the plasma membrane. New Phytol 199, 74–88.

28. Xing, S., Mehlhorn, D. G., Wallmeroth, N., Asseck, L. Y., Kar, R., Voss, A., Denninger, P., Schmidt, V. A. F., Schwarzländer, M., Stierhof, Y.-D., et al. (2017). Loss of GET pathway orthologs in Arabidopsis thaliana causes root hair growth defects and affects SNARE abundance. Proc Natl Acad Sci USA 114, E1544–E1553.

29. Zhao, Y., Araki, S., Wu, J., Teramoto, T., Chang, Y.-F., Nakano, M., Abdelfattah, A. S., Fujiwara, M., Ishihara, T., Nagai, T., et al. (2011). An expanded palette of genetically encoded Ca^2+^ indicators. Science 333, 1888–1891.

30. Keinath, N. F., Waadt, R., Brugman, R., Schroeder, J. I., Grossmann, G., Schumacher, K., and Krebs, M. (2015). Live Cell Imaging with R-GECO1 Sheds Light on flg22- and Chitin-Induced Transient [Ca^2+^]_cyt_ Patterns in Arabidopsis. Mol Plant 8, 1188–1200.

31. Simon, M. L. A., Platre, M. P., Assil, S., van Wijk, R., Chen, W. Y., Chory, J., Dreux, M., Munnik, T., and Jaillais, Y. (2014). A multi-colour/multi-affinity marker set to visualize phosphoinositide dynamics in Arabidopsis. Plant J 77, 322–337.

32. Duan, Q., Kita, D., Li, C., Cheung, A. Y., and Wu, H.-M. (2010). FERONIA receptor-like kinase regulates RHO GTPase signaling of root hair development. Proc Natl Acad Sci USA 107, 17821–17826.

33. Lin, D., Nagawa, S., Chen, J., Cao, L., Chen, X., Xu, T., Li, H., Dhonukshe, P., Yamamuro, C., Friml, J., et al. (2012). A ROP GTPase-dependent auxin signaling pathway regulates the subcellular distribution of PIN2 in Arabidopsis roots. Curr Biol 22, 1319–1325.

34. Jeon, B. W., Hwang, J. U., Hwang, Y., Song, W. Y., Fu, Y., Gu, Y., Bao, F., Cho, D., Kwak, J. M., Yang, Z., et al. (2008). The Arabidopsis Small G Protein ROP2 Is Activated by Light in Guard Cells and Inhibits Light-Induced Stomatal Opening. Plant Cell 20, 75–87.

35. Xu, T., Wen, M., Nagawa, S., Fu, Y., Chen, J.-G., Wu, M.-J., Perrot-Rechenmann, C., Friml, J., Jones, A. M., and Yang, Z. (2010). Cell surface- and rho GTPase-based auxin signaling controls cellular interdigitation in Arabidopsis. Cell 143, 99–110.

36. Fu, Y., Gu, Y., Zheng, Z., Wasteneys, G., and Yang, Z. (2005). Arabidopsis interdigitating cell growth requires two antagonistic pathways with opposing action on cell morphogenesis. Cell 120, 687–700.

37. Li, H., Lin, Y., Heath, R. M., Zhu, M. X., and Yang, Z. (1999). Control of pollen tube tip growth by a Rop GTPase-dependent pathway that leads to tip-localized calcium influx. Plant Cell 11, 1731–1742.

38. Bourne, H. R., Sanders, D. A., and McCormick, F. (1990). The GTPase superfamily: a conserved switch for diverse cell functions. Nature 348, 125–132.

39. Hodge, R. G., and Ridley, A. J. (2016). Regulating Rho GTPases and their regulators. Nat Rev Mol Cell Biol 17, 496–510.

40. Hwang, J.-U., Wu, G., Yan, A., Lee, Y.-J., Grierson, C. S., and Yang, Z. (2010). Pollen-tube tip growth requires a balance of lateral propagation and global inhibition of Rho-family GTPase activity. J Cell Sci 123, 340–350.

41. Schmidt, A., Lv, Z., and Großhans, J. (2018). ELMO and Sponge specify subapical restriction of Canoe and formation of the subapical domain in early Drosophila embryos. Development 145, dev157909.

42. Berken, A., Thomas, C., and Wittinghofer, A. (2005). A new family of RhoGEFs activates the Rop molecular switch in plants. Nature 436, 1176–1180.

43. Gu, Y., Li, S., Lord, E. M., and Yang, Z. (2006). Members of a novel class of Arabidopsis Rho guanine nucleotide exchange factors control Rho GTPase-dependent polar growth. Plant Cell 18, 366–381.

44. Qiu, J.-L., Jilk, R., Marks, M. D., and Szymanski, D. B. (2002). The Arabidopsis SPIKE1 Gene Is Required for Normal Cell Shape Control and Tissue Development. Plant Cell 14, 101–118.

45. Gu, Y., Li, S., Lord, E. M., and Yang, Z. (2006). Members of a Novel Class of Arabidopsis Rho Guanine Nucleotide Exchange Factors Control Rho GTPase-Dependent Polar Growth. Plant Cell 18, 366–381.

46. Guan, Y., Guo, J., Li, H., and Yang, Z. (2013). Signaling in Pollen Tube Growth: Crosstalk, Feedback, and Missing Links. Mol Plant 6, 1053–1064.

47. Yanagisawa, M., Alonso, J. M., and Szymanski, D. B. (2018). Microtubule-Dependent Confinement of a Cell Signaling and Actin Polymerization Control Module Regulates Polarized Cell Growth. Curr Biol 28, 2459–2466.e4.

48. Birnbaum, K., Shasha, D. E., Wang, J. Y., Jung, J. W., Lambert, G. M., Galbraith, D. W., and Benfey, P. N. (2003). A gene expression map of the Arabidopsis root. Science 302, 1956–1960.

49. Brady, S. M., Orlando, D. A., Lee, J.-Y., Wang, J. Y., Koch, J., Dinneny, J. R., Mace, D., Ohler, U., and Benfey, P. N. (2007). A high-resolution root spatiotemporal map reveals dominant expression patterns. Science 318, 801–806.

50. Borges, F., Gomes, G., Gardner, R., Moreno, N., McCormick, S., Feijó, J. A., and Becker, J. D. (2008). Comparative Transcriptomics of Arabidopsis Sperm Cells. Plant Physiol 148, 1168–1181.

51. Grossmann, G., Guo, W.-J., Ehrhardt, D. W., Frommer, W. B., Sit, R. V., Quake, S. R., and Meier, M. (2011). The RootChip: an integrated microfluidic chip for plant science. Plant Cell 23, 4234–4240.

52. Jones, A. M., Danielson, J. Å., Manojkumar, S. N., Lanquar, V., Grossmann, G., and Frommer, W. B. (2014). Abscisic acid dynamics in roots detected with genetically encoded FRET sensors. eLife 3, e01741.

53. Stanley, C. E., Shrivastava, J., Brugman, R., Heinzelmann, E., van Swaay, D., and Grossmann, G. (2018). Dual-flow-RootChip reveals local adaptations of roots towards environmental asymmetry at the physiological and genetic levels. New Phytol 217, 1357–1369.

54. Kang, E., Zheng, M., Zhang, Y., Yuan, M., Yalovsky, S., Zhu, L., and Fu, Y. (2017). The Microtubule-Associated Protein MAP18 Affects ROP2 GTPase Activity during Root Hair Growth. Plant Physiol 174, 202–222.

55. Thomas, C., Fricke, I., Weyand, M., and Berken, A. (2009). 3D structure of a binary ROP-PRONE complex: the final intermediate for a complete set of molecular snapshots of the RopGEF reaction. Biol Chem 390, 427–435.

56. Thomas, C., Fricke, I., Scrima, A., Berken, A., and Wittinghofer, A. (2007). Structural evidence for a common intermediate in small G protein-GEF reactions. Mol Cell 25, 141–149.

57. Cherfils, J., and Chardin, P. (1999). GEFs: structural basis for their activation of small GTP-binding proteins. Trends Biochem Sci 24, 306–311.

58. Wittinghofer, F. (1998). Ras signalling - Caught in the act of the switch-on. Nature News 394, 317-.

59. Li, H., Shen, J. J., Zheng, Z. L., Lin, Y., and Yang, Z. (2001). The Rop GTPase switch controls multiple developmental processes in Arabidopsis. Plant Physiol 126, 670–684.

60. Grefen, C., and Blatt, M. (2012). A 2in1 cloning system enables ratiometric bimolecular fluorescence complementation (rBiFC). Biotechniques 53.

61. Grefen, C., Obrdlik, P., and Harter, K. (2008). The Determination of Protein-protein Interactions by the Mating-based Split-ubiquitin system (mbSUS). Methods Mol Biol 479, 1–17.

62. Huang, G. Q., Li, E., Ge, F.-R., Li, S., Wang, Q., Zhang, C. Q., and Zhang, Y. (2013). Arabidopsis RopGEF4 and RopGEF10 are important for FERONIA-mediated developmental but not environmental regulation of root hair growth. New Phytol 200, 1089–1101.

63. Payne, R. J. H., and Grierson, C. S. (2009). A theoretical model for ROP localisation by auxin in Arabidopsis root hair cells. PLoS ONE 4, e8337.

64. Nagashima, Y., Tsugawa, S., Mochizuki, A., Sasaki, T., Fukuda, H., and Oda, Y. (2018). A Rho-based reaction-diffusion system governs cell wall patterning in metaxylem vessels. Sci. Rep. 8, 11542.

65. Oda, Y., Iida, Y., Kondo, Y., and Fukuda, H. (2010). Wood Cell-Wall Structure Requires Local 2D-Microtubule Disassembly by a Novel Plasma Membrane-Anchored Protein. Curr Opin Plant Biol 20, 1197–1202.

66. Bonello, T. T., Perez-Vale, K. Z., Sumigray, K. D., and Peifer, M. (2018). Rap1 acts via multiple mechanisms to position Canoe and adherens junctions and mediate apical-basal polarity establishment. Development 145, dev157941.

67. Peterson, J., Zheng, Y., Bender, L., Myers, A., Cerione, R., and Bender, A. (1994). Interactions between the bud emergence proteins Bem1p and Bem2p and Rho-type GTPases in yeast. J Cell Biol 127, 1395–1406.

68. Nern, A., and Arkowitz, R. A. (1998). A GTP-exchange factor required for cell orientation. Nature 391, 195–198.

69. Zheng, Y., Bender, A., and Cerione, R. A. (1995). Interactions among proteins involved in bud-site selection and bud-site assembly in Saccharomyces cerevisiae. J Biol Chem 270, 626–630.

70. Roemer, T., Madden, K., Chang, J., and Snyder, M. (1996). Selection of axial growth sites in yeast requires Axl2p, a novel plasma membrane glycoprotein. Genes & Development 10, 777–793.

71. Chant, J., Corrado, K., Pringle, J. R., and Herskowitz, I. (1991). Yeast BUD5, encoding a putative GDP-GTP exchange factor, is necessary for bud site selection and interacts with bud formation gene BEM1. Cell 65, 1213–1224.

72. Bender, A., and Pringle, J. R. (1989). Multicopy suppression of the cdc24 budding defect in yeast by CDC42 and three newly identified genes including the ras-related gene RSR1. Proc Natl Acad Sci USA 86, 9976–9980.

73. Chiou, J.-G., Balasubramanian, M. K., and Lew, D. J. (2017). Cell Polarity in Yeast. Annu Rev Cell Dev Biol 33, annurev–cellbio–100616–060856.

74. Wedlich-Soldner, R., Wai, S. C., Schmidt, T., and Li, R. (2004). Robust cell polarity is a dynamic state established by coupling transport and GTPase signaling. J Cell Biol 166, 889–900.

75. Yamaguchi, Y., Ota, K., and Ito, T. (2007). A Novel Cdc42-interacting Domain of the Yeast Polarity Establishment Protein Bem1 IMPLICATIONS FOR MODULATION OF MATING PHEROMONE SIGNALING. J Biol Chem 282, 29–38.

76. Butty, A. C., Pryciak, P. M., Huang, L. S., Herskowitz, I., and Peter, M. (1998). The role of Far1p in linking the heterotrimeric G protein to polarity establishment proteins during yeast mating. Science 282, 1511–1516.

77. Escobar-Restrepo, J.-M., Huck, N., Kessler, S., Gagliardini, V., Gheyselinck, J., Yang, W.-C., and Grossniklaus, U. (2007). The FERONIA receptor-like kinase mediates male-female interactions during pollen tube reception. Science 317, 656–660.

78. Hruz, T., Laule, O., Szabó, G., Wessendorp, F., Bleuler, S., Oertle, L., Widmayer, P., Gruissem, W., and Zimmermann, P. (2008). Genevestigator v3: a reference expression database for the meta-analysis of transcriptomes. Adv Bioinformatics 2008, 420747.

79. Lampropoulos, A., Sutikovic, Z., Wenzl, C., Maegele, I., Lohmann, J. U., and Forner, J. (2013). GreenGate---a novel, versatile, and efficient cloning system for plant transgenesis. PLoS ONE 8, e83043.

80. Griesbeck, O., Griesbeck, O., Baird, G. S., Baird, G. S., Campbell, R. E., Campbell, R. E., Zacharias, D. A., Zacharias, D. A., Tsien, R. Y., and Tsien, R. Y. (2001). Reducing the Environmental Sensitivity of Yellow Fluorescent Protein. J Biol Chem 276, 29188–29194.

81. Goedhart, J., Stetten, von, D., Noirclerc-Savoye, M., Lelimousin, M., Joosen, L., Hink, M. A., van Weeren, L., Gadella, T. W. J., and Royant, A. (2012). Structure-guided evolution of cyan fluorescent proteins towards a quantum yield of 93%. Nat Commun 3, 509.

82. Heim, N., Garaschuk, O., Friedrich, M. W., Mank, M., Milos, R. I., Kovalchuk, Y., Konnerth, A., and Griesbeck, O. (2007). Improved calcium imaging in transgenic mice expressing a troponin C–based biosensor. Nat Meth 4, 127–129.

83. Zuo, J., Niu, Q.-W., and Chua, N.-H. (2000). An estrogen receptor-based transactivator XVE mediates highly inducible gene expression in transgenic plants. Plant J 24, 265–273.

84. Xing, S., Wallmeroth, N., Berendzen, K. W., and Grefen, C. (2016). Techniques for the Analysis of Protein-Protein Interactions in Vivo. Plant Physiol 171, 727–758.

85. Mehlhorn, D. G., Wallmeroth, N., Berendzen, K. W., and Grefen, C. (2018). 2in1 Vectors Improve In Planta BiFC and FRET Analyses. In The Plant Endoplasmic Reticulum Methods in Molecular Biology. (New York, NY: Humana Press, New York, NY), pp. 139–158.

86. Asseck, L. Y., Wallmeroth, N., and Grefen, C. (2018). ER Membrane Protein Interactions Using the Split-Ubiquitin System (SUS). In The Plant Endoplasmic Reticulum Methods in Molecular Biology. (New York, NY: Humana Press, New York, NY), pp. 191–203.

87. Grossmann, G., Meier, M., Cartwright, H. N., Sosso, D., Quake, S. R., Ehrhardt, D. W., and Frommer, W. B. (2012). Time-lapse fluorescence imaging of Arabidopsis root growth with rapid manipulation of the root environment using the RootChip. J Vis Exp 65, e4290.

88. Stanley, C. E., Shrivastava, J., Brugman, R., Heinzelmann, E., Frajs, V., Bühler, A., van Swaay, D., and Grossmann, G. (2018). Fabrication and Use of the Dual-Flow-RootChip for the Imaging of Arabidopsis Roots in Asymmetric Microenvironments. Bio-protocol 8, 1–19.

89. Edelstein, A., Amodaj, N., Hoover, K., Vale, R., and Stuurman, N. (2010). Computer control of microscopes using μManager. Curr Protoc Mol Biol Chapter 14, Unit14.20.

90. Schindelin, J., Arganda-Carreras, I., Frise, E., Kaynig, V., Longair, M., Pietzsch, T., Preibisch, S., Rueden, C., Saalfeld, S., Schmid, B., et al. (2012). Fiji: an open-source platform for biological-image analysis. Nat Meth 9, 676–682.

