## Supplemental Material for "Distinct RopGEFs successively drive polarization and outgrowth of root hairs"

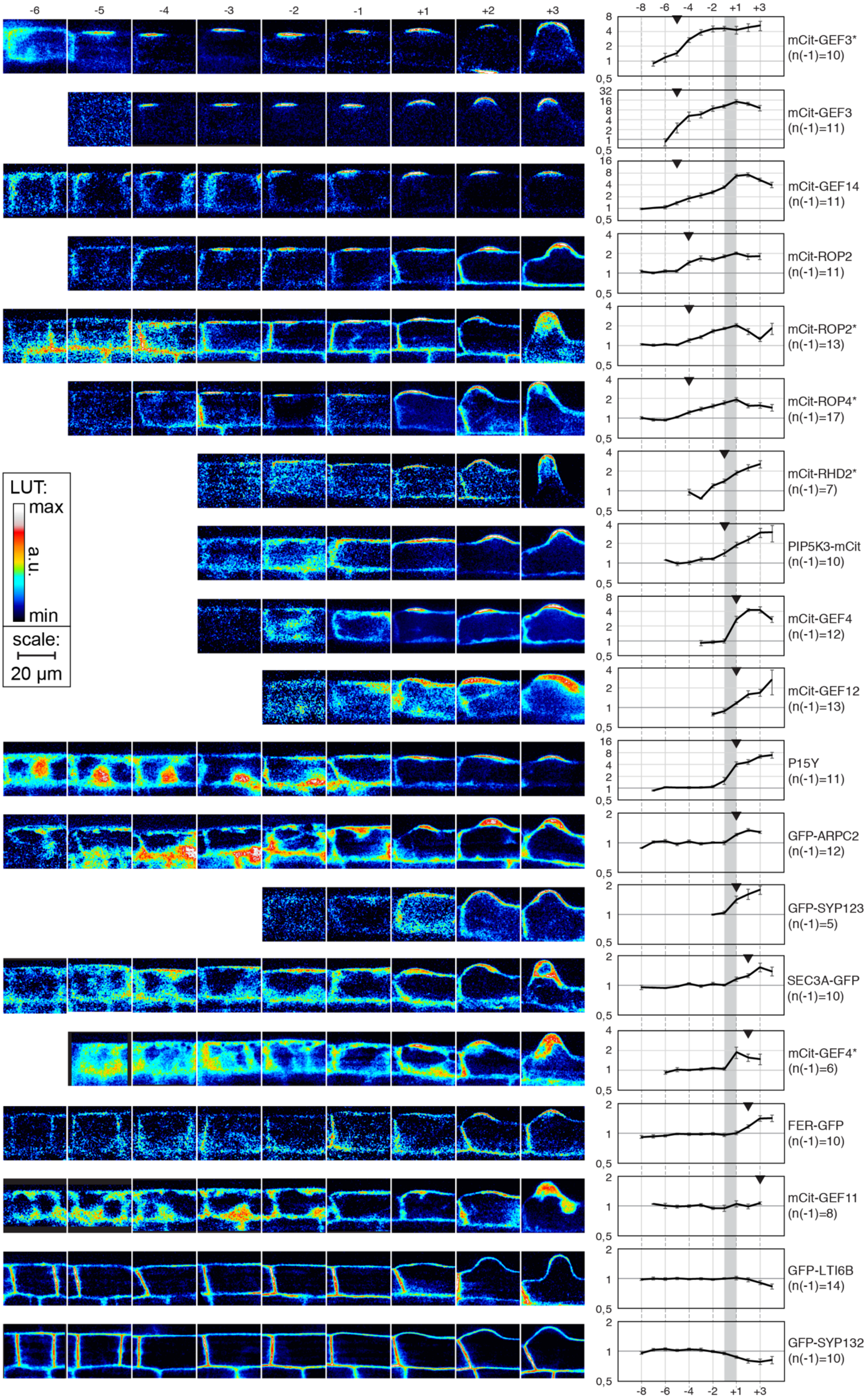

-6
-5
-4
-3
-2
-1
+1
+2
+3
LUT:
max
a.u.
min
scale:
20 µm
-8
-6
-4
-2
+1
+3
mCit-GEF3*
(n(-1)=10)
mCit-GEF3
(n(-1)=11)
mCit-GEF14
(n(-1)=11)
mCit-ROP2
(n(-1)=11)
mCit-ROP2*
(n(-1)=13)
mCit-ROP4*
(n(-1)=17)
mCit-RHD2*
(n(-1)=7)
PIP5K3-mCit
(n(-1)=10)
mCit-GEF4
(n(-1)=12)
mCit-GEF12
(n(-1)=13)
P15Y
(n(-1)=11)
GFP-ARPC2
(n(-1)=12)
GFP-SYP123
(n(-1)=5)
SEC3A-GFP
(n(-1)=10)
mCit-GEF4*
(n(-1)=6)
FER-GFP
(n(-1)=10)
mCit-GEF11
(n(-1)=8)
GFP-LTI6B
(n(-1)=14)
GFP-SYP132
(n(-1)=10)

**Figure S1. Timeline analysis of RHID markers. Related to Figure 1.**

Quantification of protein accumulation at the RHID throughout the early development of root hairs in 18 different marker lines shown in **Figure 1** & **Figure 2**. The evenly distributed membrane protein Lti6b is used as reference for protein accumulation. On the left, representative cells are shown from 6 cells before bulging (-6), to 3 cells after bulging (+3). Images are shown in intensity coding false color to highlight intensity differences. On the right, blots of the average polarity index (signal intensity in/outside RHID) for each developmental step. Error bars indicate SEM; y-axis is in $log_2$-scale; arrowheads indicate first polarity index significantly greater than reference polarity index of GFP-LTI6B (students t-test, $p < 0.05$); Grey bar indicates transition to bulging between cell -1 and +1. Corresponding genes are listed on the very right. Genes were under control of their own promoter, the *UBI10*-Promoter (P15Y), or induced by estradiol (*). Markers are sorted from earliest accumulation (GEF3*, top) to latest accumulation (GEF11, bottom). No accumulation was found for GFP-LTI6B and GFP-SYP132.

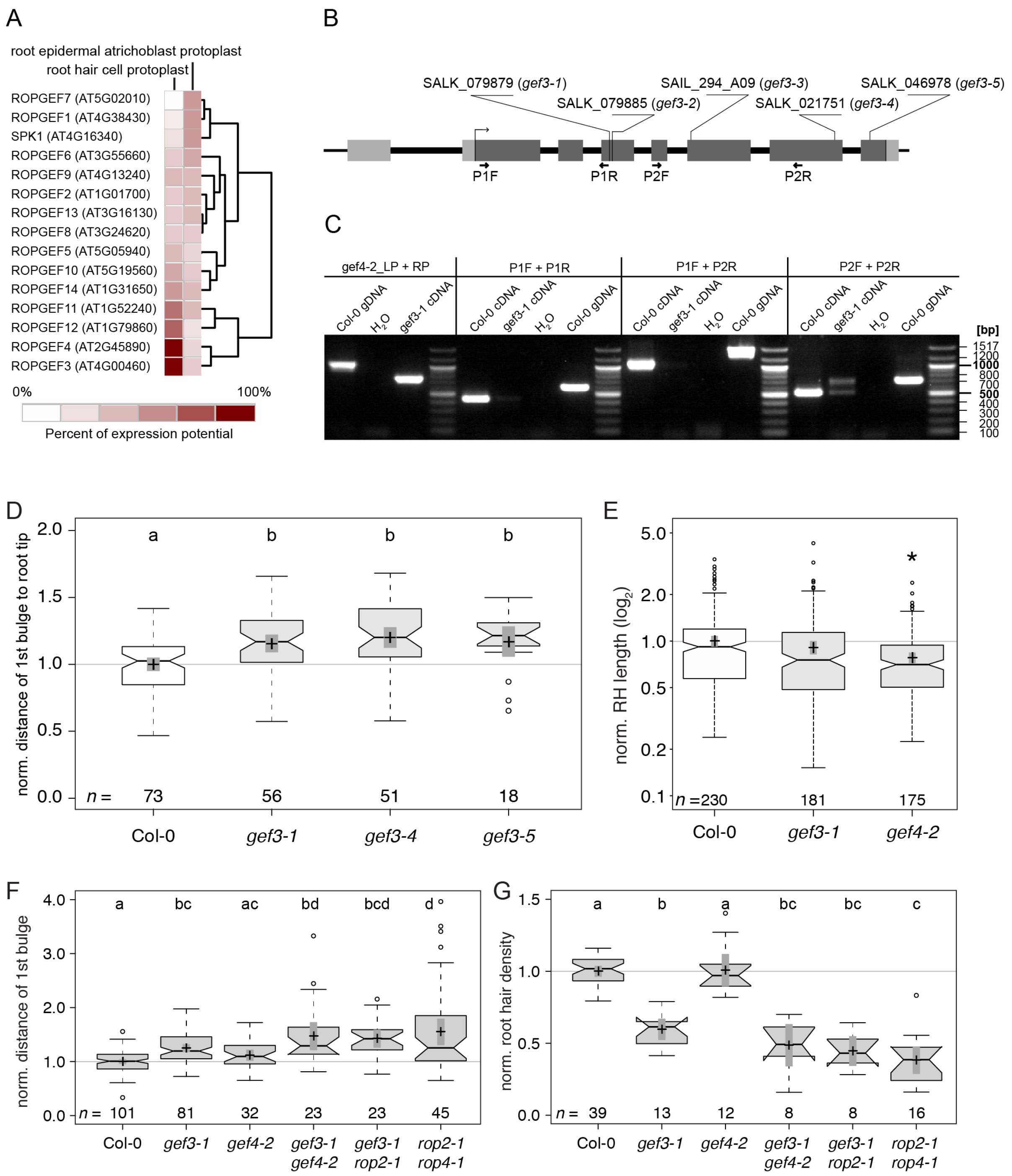

**Figure S2. Expression analysis of all *ROPGEF*s and SPIKE1, gene structure of *GEF3,* and phenotypes of single and double mutants. Related to Figure 2.**

(**A**) Comparative expression analysis of all *ROPGEF*s and SPIKE1 in root epidermis cells using hierarchical clustering of published gene expression data in GENEVESTIGATOR. Expression in trichoblasts is compared to atrichoblasts and Genes are sorted from low to high comparative trichoblast expression levels. (**B**) Gene structure of *GEF3* (AT4G00460.2) according to The Arabidopsis Information Resource

(TAIR). Black boxes show introns, grey boxes show exons (light grey: 5' and 3' UTR, dark grey translated regions). Position and ID of used T-DNA insertion lines are indicated. *gef3-1*, *gef3-4* and *gef3-5* were used for phenotyping analysis and further experiments. No homozygous plants were found for *gef3-2* and *gef3-3* in the seeds received from NASC, nor in the following generation. Those lines were not further investigated. Position of primers used for expression analysis of *GEF3* in the *gef3-1* background are indicated by black arrows. (**C**) Expression analysis of *GEF3* in Col-0 and *gef3-1*. cDNA from seedling roots (10 dag) was used. As a quality control of the *gef3-1* cDNA, *GEF4* was amplified, as it has a similar specific expression pattern compared to *GEF3*. gDNA served as a control for the PCR to exclude gDNA contamination in the samples. $H_2O$ control did not contain any PCR template. Band sizes of ladder are indicated on the right. Three primer sets for *GEF3* were used which amplified fragments in front (P1F+P1R), over (P1F+P2R) and behind (P2F+P2R) the t-DNA insertion site. (**D**) Normalized distance of first bulge to root tip measured in the indicated mutant (*gef3-4*, *gef3-5*) as compared to Col-0 and *gef3-1*. Letters show results of an ANOVA-Test (significance value 0.01), with same letters indicating no significant differences. (**E**) Quantification of hair length 3-6 mm away from root tip in *gef3-1* and gef4-2 mutant lines compared to Col-0. Normalized values are shown in $\log_2$ scale to account for the high variability in individual hairs. Asterisk indicates $p \leq 0.05$ according to two-tailed students t-test. (**F,G**) Re-plotted data on normalized distance and normalized root hair density for selected mutants for direct comparison of phenotypes. The ANOVA-Test was performed again for this set of mutants; test results are shown as letters. For a detailed explanation of all shown box-plots, see the description given in the legend of Figure 2.

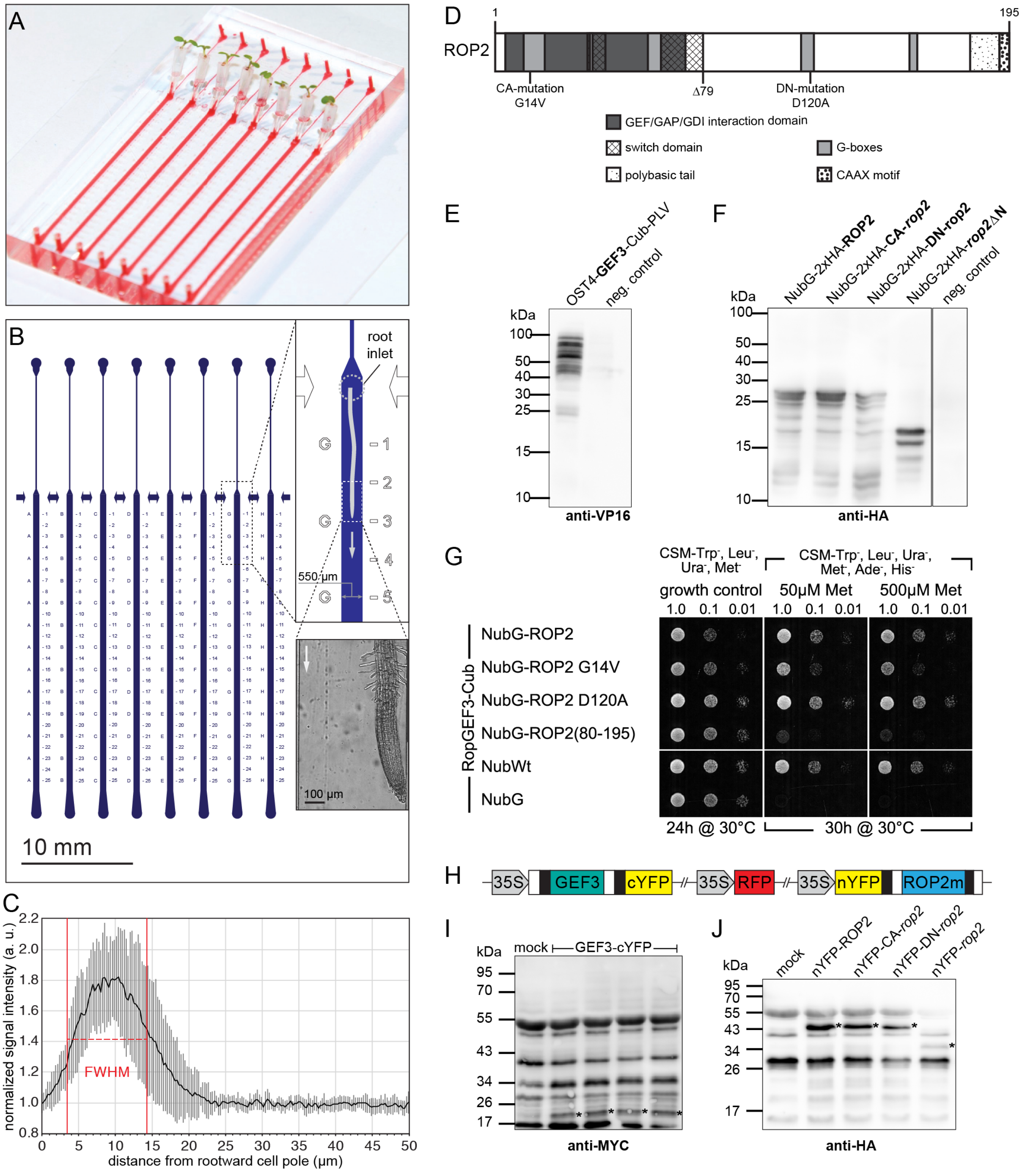

**Figure S3. The RootChip-8S, size of the polar GEF3 domain, and physical interaction between GEF3 and ROP2. Related to Figure 3.**

(**A**) Photograph of the RootChip-8S. Channels for sample perfusion are filled with red food color solution to highlight the medium channels. Arabidopsis seedlings grow through medium-filled cones into the perfusion chamber, where the root is imaged from below. (**B**) Channel design of the RootChip-8S with 8 parallel perfusion chambers and single medium inlets for each perfusion chamber (blue). Seedling inlets are marked by arrows at the top of the perfusion chambers. Labels next to each chamber help identifying the chamber (letters) and indicate the distance from the root inlet (numbers). The magnified panels show a path of a root (top right, grey sketch; bottom right, brightfield image) grown into the chamber. Grey arrows depict the direction of both medium flow and root growth. Chamber dimensions are 28 x 0.55 x 0.117 (± 0.002) mm (LxWxH). (**C**) Dimensions of the mCit-GEF3 domain at the RHID. Average intensity profile (line width 3 px) along the first 50 µm of the outer cell periphery of 17 trichoblasts just before bulging (stage -1) in GEF3::mCit-GEF3 roots. Starting position was the rootward cell border and values are normalized to the average intensity between 25 µm and 35 µm. Full width at half maximum (FWHM) of the GEF3-patch (9.6 µm ± 2.5 µm) is indicated by a dashed line. Error bars indicate standard deviation. (**D**) Schematic protein structure of ROP2. Conserved domains and motives according to NCBI Conserved Domains are indicated. The GEF/GAP/GDI interaction domain facilitates binding to regulatory proteins. The two switch domains undergo conformational changes upon regulator binding/activity status. The five G-Boxes directly bind GDP/GTP. The polybasic tail helps membrane binding and the CaaX motif is prenylated to facilitate membrane anchorage. The position of the mutations to render ROP2 constitutive active (CA-*rop2*) or dominant negative (DN-*rop2*), as well as a deletion at position 79 (*rop2*ΔN) to create a mutational construct unable to bind any effectors, are indicated. (**E**, **F**) Western blots as expression controls for bait (**E**) and prey (**F**) fusion proteins in Split-Ubiquitin-Assay. Asterisks above bands indicate full length constructs. Negative control shows protein extracts of yeast culture transformed with the corresponding 'empty' vector. (**G**) Split-ubiquitin-based assay to test for protein-protein interactions of GEF3-Cub with wildtype ROP2, CA-*rop2,* DN-*rop2* and *rop2*ΔN *(*Δ1-79), respectively. Wildtype NubI (NubWT) served as positive and NubG as negative control. (**H**) Construct used for rBiFC interaction assay. Fusion proteins and the ratiometric marker protein (RFP) are expressed from the same plasmid. GEF3 was C-terminally fused to cYFP and different versions of ROP2 N-terminally tagged with nYFP. (**I**, **J**) Western blots confirming expression of fusion proteins in rBiFC analysis. Extracts of leaves infiltrated with non-agrobacterium containing medium show cross-reactivity of antibodies with plant proteins ('Mock'). In (**I**) asterisks indicate degradation product of GEF3-cYFP. The full-length product could not be detected as this is most likely masked by the background band at ~ 55kDa. In (**J**) asterisks indicate full length constructs.

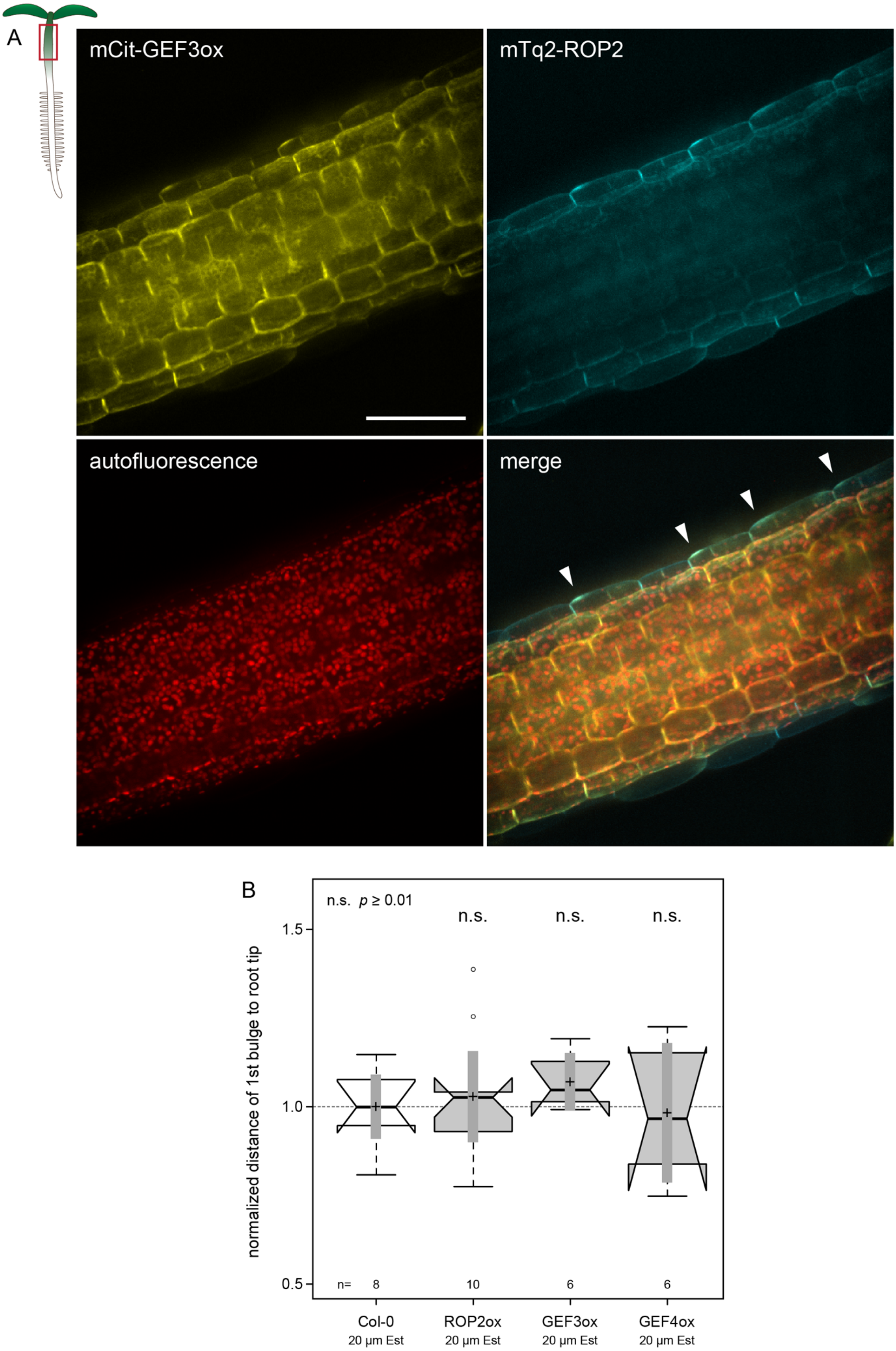

A
mCit-GEF3ox
mTq2-ROP2
autofluorescence
merge
B
n.s. $p \geq 0.01$
n.s.
n.s.
n.s.
1.5
1.0
0.5
normalized distance of 1st bulge to root tip
n=
8
10
6
6
Col-0
20 µm Est
ROP2ox
20 µm Est
GEF3ox
20 µm Est
GEF4ox
20 µm Est

**Figure S4. Ectopic, GEF3-induced polar domains in the hypocotyl epidermis, and analysis of the timing of hair emergence upon mCit-GEF3 overexpression. Related to Figure 4.**

(**A**) Overview representations of ectopic RHID-like domain formation upon induced overexpression of mCit-GEF3ox in the epidermis of the hypocotyl. Polarized mCit signal was occasionally observed (arrow heads), which then frequently coincided with ectopic accumulation of mTq2-ROP2 (expression under the control of the endogenous promoter). In 16 independent seedlings, out of 35 cells with sufficient signal of both mCit-GEF3 and mTq2-ROP2, 30 cells showed a detectable co-accumulation of both markers, five cells showed only accumulation of mCit signal. (**B**) Normalized distance of first bulge to root tip measured in Col-0 and overexpression lines of mCit-*ROP2*, mCit-*GEF3* and mCit-*GEF4* (24h after Estradiol induction). Values were normalized to corresponding Col-0 values for each independent experiment. For a detailed explanation of all shown box-plots, see the description given in the legend of Figure 2. The value of significance (*p*) was determined by a two tailed students t-test (n. s.= not significantly different).

### SUPPLEMENTAL REFERENCE

S1. Fu, Y., Gu, Y., Zheng, Z., Wasteneys, G., and Yang, Z. (2005). Arabidopsis interdigitating cell growth requires two antagonistic pathways with opposing action on cell morphogenesis. Cell *120*, 687–700.

| Gene | Gene-ID | Mutant allele | Ecotype | Mutant ID | Mutant NASC ID |
|---|---|---|---|---|---|
| ARPC2A | AT1G30825 | | | | |
| FERONIA | AT3G51550 | | | | |
| GEF3 | AT4G00460 | *gef3-1* | Col-0 | SALK_079879C | N868983 |
| | | *gef3-2* | Col-0 | SALK_079885 | N579885 |
| | | *gef3-3* | Col-0 | SAIL_294_A09 | N813628 |
| | | *gef3-4* | Col-0 | SALK_021751 | N868675 |
| | | *gef3-5* | Col-0 | SALK_046978C | N662351 |
| GEF4 | AT2G45890 | *gef4-2* | Col-0 | SALK_107520 | N607520 |
| GEF10 | AT5G19560 | *gef10-1* | Col-0 | SALK_009456 | N653017 |
| GEF11 | AT1G52240 | *gef11-1* | Col-0 | SALK_126725C | N663923 |
| GEF12 | AT1G79860 | | | | |
| GEF14 | AT1G31650 | *gef14-2* | Col-0 | SALK_046067 | N546067 |
| LTI6b | AT3G05890 | | | | |
| PCaP2 | AT5G44610 | | | | |
| PIP5K3 | AT2G26420 | | | | |
| RHD2 | AT5G51060 | | | | |
| ROP2 | AT1G20090 | *rop2-1* | Col-0 | SALK_055328C | N675195 |
| ROP4 | AT1G75840 | *rop4-1* | WS | Wisconsin T-DNA line described by Fu et al. [S1]. | |
| ROP6 | AT4G35020 | | | | |
| Sec3a | AT1G47550 | | | | |
| SYP123 | AT4G03330 | | | | |
| SYP132 | AT5G08080 | | | | |

**Table S1.** List of identifiers for genes and single mutant alleles used in this study.

### GreenGate Cloning

Double Underlined: BsaI-Sites with standart GreenGate-Overhangs;
Underlined: BsaI-Sites for fusion of multiple PCR-Fragments into one Vector;
Bold: Bases different to template for mutation

| | | | | |
|---|---|---|---|---|
| Cloning of standart modules for GrenGate cloning | Cloning of base entry vector with AD-overhangs | | agtgaagcttGGTCTCaACCTccggat<br>gcgagaattcGGTCTCaCTGAggtaccac | oPD0001-fwd<br>oPD0001D-rev |
| | Estradiol inducible Promoter module into pGGA | Ubi10-XVE with Ubi10-Terminator (reverse orientation) | aacaGGTCTCtACCTgcaacaacgaatgttcatcatactc<br>aacaGGTCTCaCTGAtctggttcgatggctgttctc | oPD0120-fwd<br>oPD0120-rev |
| | | oLexA-minimal35S-Promoter element with Linker | aacaGGTCTCaTCAGcgatcgcaccaggtaccac<br>aacaGGTCTCaTGTTcagcgtgtcctctccaaatgaa | oPD0121-fwd<br>oPD0121-rev |
| | mCitrine or mTurquoise2 into pGGC, including GAGAGA-Linker | | aaaaGGTCTCaTCAGGAGCAGGGGCGGGTGCCATGGTGAGCAAGGGCGAG<br>aaaaGGTCTCaGCAGTTACTTGTACAGCTCGTCCA | oPD0002-fwd<br>oPD0002-rev |
| | mCitrine or mTurquoise2 into pGGB, including GAGAGA-Linker | | aaaaGGTCTCaAACAATGGTGAGCAAGGGCGAGGA<br>aaaaGGTCTCaAGCCCCCGCTCCTGCTCCCTTGTACAGCTCGTCCATGCC | oPD0012-fwd<br>oPD0012-rev |
| | HSP18.2 Terminator into pGGE | | aacaGGTCTCtCTGCatatgaagatgaagatgaaatatttgg<br>aacaGGTCTCaTAGTcttatctttaatcatattccatagtcc | oPD0064-fwd<br>oPD0064-rev |

| | | | | |
|---|---|---|---|---|
| -terminal tagging only) | GEF3 Promoter with GG-Overhang A<br>GEF3 Promoter with GG-Overhang B | | aacaGGTCTCtACCTctcgatcaaccctttcacggt<br>aacaGGTCTCaTGTTctttaaaatcttaaaacactccttaaaggtct | oPD0190-fwd<br>oPD0190-rev |
| | GEF3 ORF | Part A with GG-Overhang C<br>Part A for BsaI-Site mutation | aacaGGTCTCaGGCTATGGAGAATTTATCGAATCCAGATGA<br>aacaGGTCTCCACTACAGAAGCTGTTGGTTTCATCTATCTGGTCTGAAAACCCTGAAGTGGT**t**TCTGAAC | oPD0191A-fwd<br>oPD0191A-rev |
| | | Part B<br>Part B with GG-Overhang D | aacaGGTCTCGTAGTGAAGCTTCTCCTTGTAACTG<br>aacaGGTCTCaCTGATTATTCACTACCTCTCATGGTTTTGT | oPD0191B-fwd<br>oPD0191B-rev |
| | GEF4 Promoter with GG-Overhang A<br>GEF4 Promoter with GG-Overhang B | | aacaGGTCTCtACCTctcaatgtgcataagctgctct<br>aacaGGTCTCaTGTTcacgattgtgtataatgatcaatgttt | oPD0067-fwd<br>oPD0067-rev |
| | GEF4 ORF with GG-Overhang C<br>GEF4 ORF with GG-Overhang D | | aacaGGTCTCtGGCTATGGAGAGTTCTTCGAATTCCGA<br>aacaGGTCTCtCTGACTAATCATCTCTGTTTCTCACTGTTCT | oPD0068-fwd<br>oPD0068-rev |

ry vectors - pGGA000 for promoters, pGGC000 for ORF/CDS and pGGAD000 for Promoter-ORF fusion (C-

| | | | |
|---|---|---|---|
| GEF10 Promoter with GG-Overhang A | | aacaGGTCTCtACCTggattgtgacattgagatctaccg | oPD0069-fwd |
| GEF10 Promoter with GG-Overhang B | | aacaGGTCTCaTGTTctttctaaaactaaaagatcttggcca | oPD0069-rev |
| GEF10 ORF | Part A with GG-Overhang C | aacaGGTCTCtGGCTATGTTCGATGGTCGGAACTCT | oPD0070A-fwd |
| | Part A | aacaGGTCTCCTGTCCCACCGCCTGACAT | oPD0070A-rev |
| | Part B for BsaI-Site mutation | aacaGGTCTCGGACAGGGGA**a**ACCTCTGCG | oPD0070B-fwd |
| | Part B | aacaGGTCTCTCGAAAGCTCCCTCTTCTCT | oPD0070B-rev |
| | Part C for BsaI-Site mutation | aacaGGTCTCTTTCGAAGTCCGAGCTGA**a**ACCATTTTGG | oPD0070C-fwd |
| | Part C with GG-Overhang D | aacaGGTCTCtCTGATCAGTGTCTGTCACTAGGGCT | oPD0070C-rev |
| GEF11 Promoter with GG-Overhang A | | aacaGGTCTCtACCTtttgattttgggttttgtttcgtcc | oPD0079-fwd |
| GEF11 Promoter with GG-Overhang B | | aacaGGTCTCaTGTTctttctctatctctctctctcaataac | oPD0079-rev |
| GEF11 ORF with GG-Overhang C | | aacaGGTCTCtGGCTATGTTGGAAGGCAAAGCAATGG | oPD0080-fwd |
| GEF11 ORF with GG-Overhang D | | aacaGGTCTCtCTGATCAGGAGTATCTTGCGGTTGG | oPD0080-rev |
| GEF12 Promoter with GG-Overhang A | | aacaGGTCTCtACCTatctccttttctctgtttttttttttttatttttcttc | oPD0139-fwd |
| GEF12 Promoter with GG-Overhang B | | aacaGGTCTCaTGTTtcctggtcccttgatggcaatagag | oPD0139-rev |
| GEF12 ORF | Part A with GG-Overhang C | aacaGGTCTCtGGCTATGGTTCGTGCTTCGGAACA | oPD0140A-fwd |
| | Part A for BsaI-Site mutation | aacaGGTCTCGATCCGGTGGCAGCCTCATCTAAACC**a**GTCTCGAC | oPD0140A-rev |
| | Part B | aacaGGTCTCCGGATCCCATGACGCTGAA | oPD0140B-fwd |
| | Part B | aacaGGTCTCTCTCGCTTCTCCAGACTTACTC | oPD0140B-rev |
| | Part C for BsaI-Site mutation | aacaGGTCTCGCGAGAGGTCTTCGAAGAGCGAGCTGA**a**ACCATTTTG | oPD0140C-fwd |
| | Part C with GG-Overhang D | aacaGGTCTCtCTGATCAATGCCGTGCCGTTGG | oPD0140C-rev |
| GEF14 Promoter with GG-Overhang A | | aacaGGTCTCtACCTGATGCATTGGTTGCTCACTTCA | oPD0141-fwd |
| GEF14 Promoter with GG-Overhang B | | aacaGGTCTCaTGTTcctttcttctcttttgaattcttttgttga | oPD0141-rev |
| GEF14 ORF | Part A with GG-Overhang C | aacaGGTCTCtGGCTATGATGCTGATGAGAAGAAGGTTG | oPD0142A-fwd |
| | Part A | aacaGGTCTCactataaagaaaaccccacctagtacat | oPD0142A-rev |
| | Part B for BsaI-Site mutation | aacaGGTCTCtatagtcaaaatctgaac**A**gtctcttaatg | oPD0142B-fwd |
| | Part B for BsaI-Site mutation | aacaGGTCTCTTAGCATCAAAATGTTGAGA**t**CTTCCAG | oPD0142B-rev |
| | Part C | aacaGGTCTCTGCTAAGGAGAAGAAGAAACAAGGC | oPD0142C-fwd |
| | Part C | aacaGGTCTCGGAGCTAGGGAAGCAACTCG | oPD0142C-rev |
| | Part D for BsaI-Site mutation | aacaGGTCTCAGCTCCCGTGACCCTTATAGGACACCTGA**a**AGACCTC | oPD0142D-fwd |
| | Part D with GG-Overhang D | aacaGGTCTCaCTGAAGGAGAAGTATCAGAAGGCACTTTAC | oPD0142D-rev |
| PCaP2 ORF with GG-Overhang C | | aacaGGTCTCaGGCTATGGGTTATTGGAAGTCGAAGGT | oPD0129-fwd |
| PCaP2 ORF with GG-Overhang D | | aaaaGGTCTCtCTGAAGCCTTTTGTGGCGCAGCCG | oPD0003-rev |
| | Part A with GG-Overhang A | aacaGGTCTCtACCTgccaatcttcttgtcatctacatcc | oPD0060A-fwd |

| | | | | |
|---|---|---|---|---|
| Cloning of Promoters and ORF/CDS into GreenGate ent[illegible] | PIP5K3 (Promoter and ORF) for cloning into pGGAD | Part A with GG-Overhang A | aacaGGTCTCCAGCCGTATCACTCACCTCG | oPD0060A-rev |
| | | Part B | aacaGGTCTCCGGCTGCCGAGATTAGAATAGT | oPD0060B-fwd |
| | | Part B for BsaI-Site mutation | aacaGGTCTCGATCCAAGTCTTTGAGTGTTGT**c**GTCTCGTCG | oPD0060B-rev |
| | | Part C for BsaI-Site mutation | aacaGGTCTCTGGATCTCAAGTATGTGTTTCGACTCGA**a**ACCTCATGG | oPD0060C-fwd |
| | | Part C for BsaI-Site mutation | aacaGGTCTCCTCGTCACGCATGCCAGATTCACGGAAATG**c**AGACCAATC | oPD0060C-rev |
| | | Part D | aacaGGTCTCGACGACATTTCCTTGGGCATC | oPD0060D-fwd |
| | | Part D with GG-Overhang D | aacaGGTCTCtCTGATTTGTCTTCAATGAATATTTTGT | oPD0060D-rev |
| | RHD2 ORF | Part A with GG-Overhang C | aacaGGTCTCaGGCTATGTCTAGAGTGAGTTTTGAAGTGT | oPD0087A-fwd |
| | | Part A | aacaGGTCTCcACACcgctttattataatgaattaggt | oPD0087A-rev |
| | | Part B for BsaI-Site mutation | aacaGGTCTCgGTGTgggagac**G**acaactaaaa | oPD0087B-fwd |
| | | Part B | aacaGGTCTCCCTCGTACTTCTTGTAGTCTTGTGC | oPD0087B-rev |
| | | Part C for BsaI-Site mutation | aacaGGTCTCACGAGGTGGTTCTACTAGTTGGTtT**a**GGGATTGG | oPD0087C-fwd |
| | | Part C with GG-Overhang D | aacaGGTCTCaCTGATTAGAAATTCTCTTTGTGGAAGGA | oPD0087C-rev |
| | ROP2 Promoter with GG-Overhang A | | aacaGGTCTCtACCTgacaaataattataagaagctaccgtctg | oPD0038-fwd |
| | ROP2 Promoter with GG-Overhang B | | aaaaGGTCTCtTGTTctctgccgcaagatcggaaa | oPD0006-rev |
| | ROP2 ORF & CDS with GG-Overhang C | | aaaaGGTCTCaGGCTATGGCGTCAAGGTTTATAAAGTGTG | oPD0007B-fwd |
| | ROP2 ORF & CDS with GG-Overhang D | | aaaaGGTCTCtCTGATCACAAGAACGCGCAACGGT | oPD0007-rev |
| | ROP4 ORF with GG-Overhang C | | aaaaGGTCTCaGGCTATGAGTGCTTCGAGGTTTAT | oPD0009-fwd |
| | ROP4 ORF with GG-Overhang D | | aaaaGGTCTCtCTGATCACAAGAACACGCAGCGGT | oPD0009-rev |
| | ROP6 ORF with GG-Overhang C | | aaaaGGTCTCaGGCTATGAGTGCTTCAAGGTTTAT | oPD0011-fwd |
| | ROP6 ORF with GG-Overhang D | | ttttGGTCTCtCTGATCAGAGTATAGAACAACCTT | oPD0011-rev |

| | | | |
|---|---|---|---|
| Mutagenesis of ROP2 | rop2$^{CA}$-mutation (G14V) with oPD0007-rev Primer | aacaGGTCTCtGGCTATGGCGTCAAGGTTTATAAAGTGTGTGACCGTCGGAGATG**t**TGCCGTCGG | oPD0092-fwd |
| | rop2$^{DN}$-mutation (D120A) with oPD0007B-fwd Primer | aacaGGTCTCTTGTCCCAACAAGGATAATGGGAA | oPD0093A-rev |
| | rop2$^{DN}$-mutation (D120A) with oPD0007-rev Primer | aacaGGTCTCGGACAAAACTCG**c**TCTTCGAGAT | oPD0093B-fwd |
| | rop2$^{\Delta20}$-mutation with oPD0007-rev Primer | aacaGGTCTCtGGCTTGCATGCTCATTTCTTACACTAGC | oPD0094-fwd |
| | rop2$^{\Delta43}$-mutation with oPD0007-rev Primer | aacaGGTCTCtGGCTAGTGCTAATGTGGTTGTTGATGG | oPD0095-fwd |
| | rop2$^{\Delta80}$-mutation with oPD0007-rev Primer | aacaGGTCTCtGGCTTTCATTCTTGCTTTCTCTCTTATTAGCA | oPD0096-fwd |

| | | | | |
|---|---|---|---|---|
| | Gef3-P1F | Expression Analysis of Gef3 | TGAAAACGACGATCATCAATCACC | oPD0231-fwd |
| | Gef3-P1R | Expression Analysis of Gef3 | GGCTCTAACCTCAGATTCTGTCC | oPD0231-rev |
| | Gef3-P2F | Expression Analysis of Gef3 | TGGAGAGTAGACCAAGAGCAGA | oPD0232-fwd |
| | Gef3-P2R | Expression Analysis of Gef3 | TCTCCATGTGTACATCGAAGCT | oPD0232-rev |

| | | | | |
|---|---|---|---|---|
| Expression test and Genotyping of t-DNA lines | gef3-1_LP<br>gef3-1_RP | Genotyping of gef3-1 (SALK_079879C) & gef3-2 (SALK_079885) | TCGAATCCAGATGAAAACGAC<br>TCCTGAATGATCCAGTCGAAG | oPD0197-LP<br>oPD0197-RP |
| | gef3-3_LP<br>gef3-3_RP | Genotyping of gef3-3 (SAIL_294_A09) | CGCTGTTTCAAACAGAAGAGG<br>CAGAATCTGAGGTTAGAGCCG | oAR012-LP<br>oAR012-RP |
| | gef3-4_LP<br>gef3-4_RP | Genotyping of gef3-3 (SALK_021751) | AGAAAGGAGCAAAAAGCTTGG<br>GATTCATAAAGCTGCAATGGC | oAR013-LP<br>oAR013-RP |
| | gef3-5_LP<br>gef3-5_RP | Genotyping of gef3-5 (SALK_046978C) | TCGAAGATGGGACAAGTTCAG<br>TGCAGTGTGGTAAAAGCAGTG | oAR014-LP<br>oAR014-RP |
| | gef4-2_LP<br>gef4-2_RP | Genotyping of gef4-2 (SALK_107520) | AACCTTCAGCAGGAACACATG<br>AGAGTTCTTCGAATTCCGACC | oPD0181-LP<br>oPD0181-RP |
| | gef10-1_LP<br>gef10-1_RP | Genotyping of gef10-1 (SALK_009456) | TTGACCGAAATAAGAAGTCCTC<br>TTTGAAACGATGTGGTGTTTG | oPD0110-LP<br>oPD0110-RP |
| | gef11-1_LP<br>gef11-1_RP | Genotyping of gef11-1 (SALK_126725C) | TCAGAGAGAGGTCAAAATTGAGG<br>TACCTGCGAGATTGGTAATGG | oPD0168-LP<br>oPD0168-RP |
| | gef14-2_LP<br>gef14-2_RP | Genotyping of gef14-2 (SALK_046067) | TGGTAAGACACCGAAACTTGC<br>TGCTGATGAGAAGAAGGTTG | oPD0176-LP<br>oPD0176-RP |
| | rop2-1_LP<br>rop2-1_RP | Genotyping of rop2-1 (SALK_055328C) | TCGAATTTGGGTGATTCTCAG<br>TGTGGACTCGAAAGATTCACC | oPD0104-LP<br>oPD0104-RP |
| | rop4-1_LP<br>rop4-1_RP | Genotyping of rop4-1 (Wisconsin t-DNA line, Fu et al. 2005, Cell) | aaaaGGTCTCaGGCTATGAGTGCTTCGAGGTTTAT<br>aaaaGGTCTCtCTGATCACAAGAACACGCAGCGGT | oPD0009-fwd<br>oPD0009-rev |
| | LB1.3x_Salk | internal Primer for Salk t-DNA lines | GGATTTTGCCGATTTCGGAACCACC | |
| | LB3_Sail | internal Primer for Sail t-DNA lines | TAGCATCTGAATTTCATAACCAATCTCGATACAC | |
| | JL-202 | internal Primer for Wisconsin t-DNA lines | CATTTTATAATAACGCTGCGGACATCTAC | |

### Gateway cloning

Small Letters: attB Site
Red: Additional bases for correct reading frame

| | | |
|---|---|---|
| Cloning of PCR fragments for Gateway cloning | DM-GEF3 -attB1 | ggggacaagtttgtacaaaaaagcaggctCTATGGAGAATTTATCGAATCCAGATG |
| | DM-GEF3 -attB2-ST | ggggaccactttgtacaagaaagctgggtTTATTCACTACCTCTCATGGTTTTG |
| | DM-GEF3 -attB2-wo | ggggaccactttgtacaagaaagctgggtgTTCACTACCTCTCATGGTTTTG |
| | DM-GEF3 -attB4-ST | ggggacaactttgtatagaaaagttgggtTTATTCACTACCTCTCATGGTTTTG |
| | DM-GEF3 -attB4-wo | ggggacaactttgtatagaaaagttgggtgTTCACTACCTCTCATGGTTTTG |
| | DM-ROP2 -attB1 | ggggacaagtttgtacaaaaaagcaggctCTATGGCGTCAAGGTTTATAAAGTGTG |
| | DM-ROP2 -attB2-ST | ggggaccactttgtacaagaaagctgggtTCACAAGAACGCGCAACGG |
| | DM-ROP2 -attB3 | ggggacaactttgtataataaagttgCTATGGCGTCAAGGTTTATAAAGTGTG |
| | DM-rop2ΔN -attB1 | ggggacaagtttgtacaaaaaagcaggctCTTTCATTCTTGCTTTCTCTCTTATTAGC |
| | DM-rop2ΔN -attB3 | ggggacaactttgtataataaagttgCTTTCATTCTTGCTTTCTCTCTTATTAGC |

**Table S2.** List of PCR primers used for GreenGate and Gateway cloning.

| | | Promoter | N-terminal Tag | ORF / CDS | C-terminal Tag | Terminator | Plant selection marker | Plant Expression Vector | Bacterial selection marker | Plasmid ID |
|---|---|---|---|---|---|---|---|---|---|---|
| | | Module A | Module B | Module C | Module D | Module E | Module F | Module Z | | |
| GreenGate expression vectors | **GEF3-mCitrine** | GEF3-Promoter | mCitrine w/ Linker | GEF3-ORF | Decoy | HSP18.2 | Basta | pGGZ003 | Spec/Strep | pPD0332 |
| | **GEF3-mCitrine (inducible)** | Ubi-XVE_oLexA-35S | mCitrine w/ Linker | GEF3-ORF | Decoy | HSP18.2 | Basta | pGGZ003 | Spec/Strep | pPD0333 |
| | **GEF3-mTurquoise2 (inducible)** | Ubi-XVE_oLexA-35S | mTurquoise2 w/ Linker | GEF3-ORF | Decoy | HSP18.2 | Kanamycin | pGGZ003 | Spec/Strep | pPD0334 |
| | **GEF4-mCitrine** | GEF4-Promoter | mCitrine w/ Linker | GEF4-ORF | Decoy | HSP18.2 | Basta | pGGZ003 | Spec/Strep | pPD0301 |
| | **GEF4-mCitrine (inducible)** | Ubi-XVE_oLexA-35S | mCitrine w/ Linker | GEF4-ORF | Decoy | HSP18.2 | Basta | pGGZ003 | Spec/Strep | pPD0202 |
| | **GEF10-mCitrine** | GEF10-Promoter | mCitrine w/ Linker | GEF10-ORF | Decoy | HSP18.2 | Basta | pGGZ003 | Spec/Strep | pPD0302 |
| | **GEF10-mCitrine (inducible)** | Ubi-XVE_oLexA-35S | mCitrine w/ Linker | GEF10-ORF | Decoy | HSP18.2 | Basta | pGGZ003 | Spec/Strep | pPD0203 |
| | **GEF11-mCitrine** | GEF11-Promoter | mCitrine w/ Linker | GEF11-ORF | Decoy | HSP18.2 | Basta | pGGZ003 | Spec/Strep | pPD0303 |
| | **GEF11-mCitrine (inducible)** | Ubi-XVE_oLexA-35S | mCitrine w/ Linker | GEF11-ORF | Decoy | HSP18.2 | Basta | pGGZ003 | Spec/Strep | pPD0204 |
| | **GEF12-mCitrine** | GEF12-Promoter | mCitrine w/ Linker | GEF12-ORF | Decoy | HSP18.2 | Basta | pGGZ003 | Spec/Strep | pPD0304 |
| | **GEF12-mCitrine (inducible)** | Ubi-XVE_oLexA-35S | mCitrine w/ Linker | GEF12-ORF | Decoy | HSP18.2 | Basta | pGGZ003 | Spec/Strep | pPD0205 |
| | **GEF14-mCitrine** | GEF14-Promoter | mCitrine w/ Linker | GEF14-ORF | Decoy | HSP18.2 | Basta | pGGZ003 | Spec/Strep | pPD0305 |
| | **GEF14-mCitrine (inducible)** | Ubi-XVE_oLexA-35S | mCitrine w/ Linker | GEF14-ORF | Decoy | HSP18.2 | Basta | pGGZ003 | Spec/Strep | pPD0206 |
| | **PCaP2-mCitrine (inducible)** | Ubi-XVE_oLexA-35S | Decoy | PCaP2-ORF | mCitrine w/ Linker | HSP18.2 | Basta | pGGZ003 | Spec/Strep | pPD0215 |
| | **PIP5K3-mCitrine** | PI5K3-Promoter and ORF | | | mCitrine w/ Linker | HSP18.2 | Basta | pGGZ003 | Spec/Strep | pPD0241 |
| | **RHD2-mCitrine (inducible)** | Ubi-XVE_oLexA-35S | mCitrine w/ Linker | RHD2-ORF | Decoy | HSP18.2 | Basta | pGGZ003 | Spec/Strep | pPD0209 |
| | **ROP2-mTurquoise2** | ROP2-Promoter | mTurquoise2 w/ Linker | ROP2-ORF | Decoy | RubisCO | Kanamycin | pGGZ003 | Spec/Strep | pPD0042 |
| | **ROP2-mCitrine** | ROP2-Promoter | mCitrine w/ Linker | ROP2-ORF | Decoy | HSP18.2 | Basta | pGGZ003 | Spec/Strep | pPD0240 |
| | **ROP2-ORF-mCitrine (inducible)** | Ubi-XVE_oLexA-35S | mCitrine w/ Linker | ROP2-ORF | Decoy | HSP18.2 | Basta | pGGZ003 | Spec/Strep | pPD0199 |
| | **ROP2-CDS-mCitrine (inducible)** | Ubi-XVE_oLexA-35S | mCitrine w/ Linker | ROP2-CDS | Decoy | HSP18.3 | Basta | pGGZ003 | Spec/Strep | pPD0184 |
| | **rop2ΔN-mCitrine (inducible)** | Ubi-XVE_oLexA-35S | mCitrine w/ Linker | rop2ΔN-CDS | Decoy | HSP18.2 | Basta | pGGZ003 | Spec/Strep | pPD0195 |
| | **ROP4-mCitrine (inducible)** | Ubi-XVE_oLexA-35S | mCitrine w/ Linker | ROP4-ORF | Decoy | HSP18.2 | Basta | pGGZ003 | Spec/Strep | pPD0200 |
| | **ROP6-mCitrine (inducible)** | Ubi-XVE_oLexA-35S | mCitrine w/ Linker | ROP6-ORF | Decoy | HSP18.2 | Basta | pGGZ003 | Spec/Strep | pPD0201 |
| Gateway expression vectors | **NubG-ROP2** | ADH | NubG w/ 2xHA | ROP2-CDS | - | - | Yeast: TRP | - | Amp | D1288 |
| | **NubG-CA-ROP2** | ADH | NubG w/ 2xHA | CA-ROP2-CDS | - | - | Yeast: TRP | - | Amp | D1289 |
| | **NubG-DN-ROP2** | ADH | NubG w/ 2xHA | DN-ROP2-CDS | - | - | Yeast: TRP | - | Amp | D1290 |
| | **NubG-rop2ΔN** | ADH | NubG w/ 2xHA | rop2ΔN-CDS | - | - | Yeast: TRP | - | Amp | D1291 |
| | **Ost4-GEF3-Cub** | met25 | mOst4 | GEF3-CDS | Cub | - | Yeast: Leu | - | Amp | D1295 |
| | **GEF3-cYFP--nYFP-ROP2** | 3x35S | nYFP-HA | GEF3-CDS/ RFP/ROP2-CDS | MYC-cYFP | T35S | - | - | Spec | D1189 |
| | **GEF3-cYFP--nYFP-CA-ROP2** | 3x35S | nYFP-HA | GEF3-CDS/RFP/ CA-ROP2-CDS | MYC-cYFP | T35S | - | - | Spec | D1190 |
| | **GEF3-cYFP--nYFP-DN-ROP2** | 3x35S | nYFP-HA | GEF3-CDS/RFP/ DN-ROP2-CDS | MYC-cYFP | T35S | - | - | Spec | D1191 |
| | **GEF3-cYFP--nYFP-rop2ΔN** | 3x35S | nYFP-HA | GEF3-CDS/RFP/ rop2ΔN-CDS | MYC-cYFP | T35S | - | - | Spec | D1192 |
| | **FER-GFP** | FER | - | FER | mGFP6 | NOS | Hygromycin | pMDC111 | Kanamycin | pMDC111 |

**Table S3**. List of used expression vectors.

```
> # This script has been modified from the script #84 TUKEY TEST, deposited on R-graph-gallery
(https://www.r-graph-gallery.com/84-tukey-test/) by Yan Holtz, Queensland Brain Institute,
University of Queensland.

> # script modifications by Guido Grossmann, Centre for Organismal Studies, Heidelberg
University
> # library
> library(multcompView)
>
> # Import root hair distance data
> data <- read.csv("~/distance.csv",header=TRUE)
>
> # What is the effect of the line on the value?
> model=lm( data$value ~ data$line )
> ANOVA=aov(model)
>
> # Tukey test to study each mutant line pair:
> TUKEY <- TukeyHSD(x=ANOVA, 'data$line', conf.level=0.99)
>
> # Tukey test representation :
> plot(TUKEY , las=1 , col="brown" )
>
> # Grouping the mutant lines that are not different to each other.
> generate_label_df <- function(TUKEY, variable){
+ # Extract labels and factor levels from Tukey post-hoc
+ Tukey.levels <- TUKEY[[variable]][,4]
+ Tukey.labels <- data.frame(multcompLetters(Tukey.levels)['Letters'])
+
+ #Ordering the labels as in the boxplot :
+ Tukey.labels$line=rownames(Tukey.labels)
+ Tukey.labels=Tukey.labels[order(Tukey.labels$line) , ]
+ return(Tukey.labels)
+ }
> # Apply the function on the dataset
> LABELS=generate_label_df(TUKEY , "data$line")
>
> # A panel of colors to draw each group with the same color :
> my_colors=c( rgb(143,199,74,maxColorValue = 255),rgb(242,104,34,maxColorValue = 255),
rgb(111,145,202,maxColorValue = 255),rgb(254,188,18,maxColorValue = 255) ,
rgb(74,132,54,maxColorValue = 255),rgb(236,33,39,maxColorValue =
255),rgb(165,103,40,maxColorValue = 255))
>
> # Draw the basic boxplot
> a=boxplot(data$value ~ data$line , ylim=c(min(data$value) , 1.1*max(data$value)) ,
col=my_colors[as.numeric(LABELS[,1])] , ylab="value" , main="")
>
> # Adding a letter over each box. Over is how high I want to write it.
> over=0.1*max( a$stats[nrow(a$stats),] )
>
> #Adding the labels
> text( c(1:nlevels(data$line)) , a$stats[nrow(a$stats),]+over , LABELS[,1]  ,
col=my_colors[as.numeric(LABELS[,1])] )
```
